# An engineered CRISPR/Cas9 mouse line for simultaneous readout of lineage histories and gene expression profiles in single cells

**DOI:** 10.1101/797597

**Authors:** Sarah Bowling, Duluxan Sritharan, Fernando G. Osorio, Maximilian Nguyen, Priscilla Cheung, Alejo Rodriguez-Fraticelli, Sachin Patel, Yuko Fujiwara, Bin E. Li, Stuart H. Orkin, Sahand Hormoz, Fernando D. Camargo

**Affiliations:** Stem Cell Program, Boston Children’s Hospital, Boston, MA, USA; Department of Stem Cell and Regenerative Biology, Harvard University, Cambridge, MA, USA; Harvard Graduate Program in Biophysics, Harvard University, Cambridge, MA, USA; Department of Data Sciences, Dana-Farber Cancer Institute, Boston, MA, USA; Department of Systems Biology, Harvard Medical School, Boston, MA, USA; Division of Hematology/Oncology, Boston Children’s Hospital, Harvard Medical School, Boston, MA, USA; Department of Pediatric Oncology, Dana-Farber Cancer Institute, Boston, MA, USA; Howard Hughes Medical Institute, Boston, MA, USA; Broad Institute of MIT and Harvard, Cambridge, MA, USA

## Abstract

Tracing the lineage history of cells is key to answering diverse and fundamental questions in biology. Particularly in the context of stem cell biology, analysis of single cell lineages in their native state has elucidated novel fates and highlighted heterogeneity of function. Coupling of such ancestry information with other molecular readouts represents an important goal in the field. Here, we describe the CARLIN (for <u>C</u>RISPR <u>A</u>rray <u>R</u>epair <u>LIN</u>eage tracing) mouse line and corresponding analysis tools that can be used to simultaneously interrogate the lineage and transcriptomic information of single cells *in vivo.* This model exploits CRISPR technology to generate up to 44,000 transcribed barcodes in an inducible fashion at any point during development or adulthood, is compatible with sequential barcoding, and is fully genetically defined. We have used CARLIN to identify intrinsic biases in the activity of fetal liver hematopoietic stem cell (HSC) clones and to uncover a previously unappreciated clonal bottleneck in the response of HSCs to injury. CARLIN also allows the unbiased identification of transcriptional signatures based on *in vivo* stem cell function without a need for markers or cell sorting.

## Introduction

Generating animal models that enable cell lineage tracing *in vivo* has been a long-standing aim in biological research. Historically, lineage tracing in mammals has been limited to labelling and tracking small populations of cells through the use of dyes or fluorescent markers (Kretzschmar and Watt, 2012). Although these techniques helped resolve major questions in biology, from lineage commitment during early development (Balakier and Pedersen, 1982) to adult stem cell behavior (Snippert et al., 2010), the low number of clones analyzed at any one timepoint limit comprehensive understanding of global stem cell dynamics within tissues. These approaches are also intrinsically limited in their ability to trace individual cells and therefore provide limited insight into heterogeneity in cell populations. Retrovirally-delivered DNA barcodes have been used as clonal markers particularly in the context of blood generation (Gerrits et al., 2010; Lu et al., 2011; Schepers et al., 2008). However, introduction of such barcodes requires stem cells to be extracted from the tissue and manipulated. Recently, two mouse models have been developed that enable barcoding of cells in their native environment using randomly integrated transposons (Sun et al., 2014) or recombinases that create genetic diversity in a distinct locus (Pei et al., 2017). The use of these models has revealed dramatic differences between hematopoietic stem cell (HSC) behavior in unperturbed conditions versus transplantation, and has highlighted important functional heterogeneity within the HSC compartment (Pei et al., 2017; Rodriguez-Fraticelli et al., 2018). However, these models are limited in that they only provide lineage information and do not provide molecular insight into the genetic program driving heterogeneous behavior.

The advent of CRISPR/Cas9 has led to the development of additional lineage tracing tools that use errors from non-homologous end-joining DNA repair to generate a high diversity of unique and heritable DNA barcodes. A first proof-of-principle study demonstrated the feasibility of lineage tracing via this method in the zebrafish embryo (McKenna et al., 2016). Modified variants of this system that use expressed barcodes have allowed for the simultaneous measurement of single-cell gene expression levels and lineage tracing (Alemany et al., 2018; Raj et al., 2018; Spanjaard et al., 2018). More recently, systems combining delivery of expressed barcodes with transposons in the mouse embryo have been described (Chan et al., 2019; Kalhor et al., 2018). Both of these approaches use constitutively expressed Cas9 and use multiple target arrays (barcodes) to generate diversity. However, new embryonic manipulations are required to generate mice every time, and the resulting mice are impractical for breeding given the high number of randomly inserted transgenes. Therefore, these models are unsuitable for lineage analysis of adult tissues.

Here we present a versatile mouse model that allows inducible CRISPR-based lineage tracing that is genetically defined, incorporates inducible, transcribed barcodes, and works across adult mouse tissues. Furthermore, we have developed the analysis tools and reference data sets required to interpret the detected barcodes and quantify their statistical significance. Due to our ability to simultaneously interrogate lineage and transcriptional profiles of single-cells, the CARLIN system presents unique advantages to study stem cell clonal dynamics compared with previously generated cell lineage tracing tools. We exploit these advantages here to unveil unknown aspects of hematopoiesis during development and in adulthood following stress.

## Results

### Inducible and dose-dependent molecular barcoding in mouse embryonic stem cells

We set out to generate a genetically-defined mouse model in which we could 1) record the lineage histories of individual cells within their own DNA and 2) read out lineage histories alongside gene expression profiles at the single-cell level. Based on the GESTALT model that has been successfully used for molecular recording in zebrafish (McKenna et al., 2016), we designed 10 sgRNAs that enable efficient cutting of target sites in the presence of Cas9 (Supplementary Figure 1A) with minimal off-target activity within the mouse genome (Methods). We designed the gRNA cassette in two iterations, one where individual U6 promoters drove sgRNA expression (Figure 1A), and a second cassette carrying tetO-operons upstream of each sgRNA (iCARLIN, Supplementary Figure 1B). Unless otherwise stated, all data presented here correspond to the first system. We also designed a 276 bp array containing target sites perfectly matching each of the expressed gRNAs (Figure 1A,B). Constitutive expression of the molecular recorder array is achieved through its placement in the 3’ UTR of a fluorescent protein driven by the constitutive CAG promoter. All of these elements were inserted together in the widely-used *Col1a1* locus via recombinase-mediated integration into mouse embryonic stem (ES) cells that also express an enhanced reverse tetracycline trans-activator (M2-rtTA) from the ubiquitous Rosa26 promoter (Beard et al., 2006). We then generated mouse lines from these ES cells. To have temporal control of Cas9 expression, we separately created a mouse line that expresses both Doxycyline-dependent Cas9 (tetO-Cas9; integrated in the *Col1a1* locus) and M2-rtTA (integrated in the *Rosa26* locus). Finally, we crossed these two mouse lines to generate CARLIN mice and CARLIN ES cells that carry all the transgenic elements.

**Figure 1:**
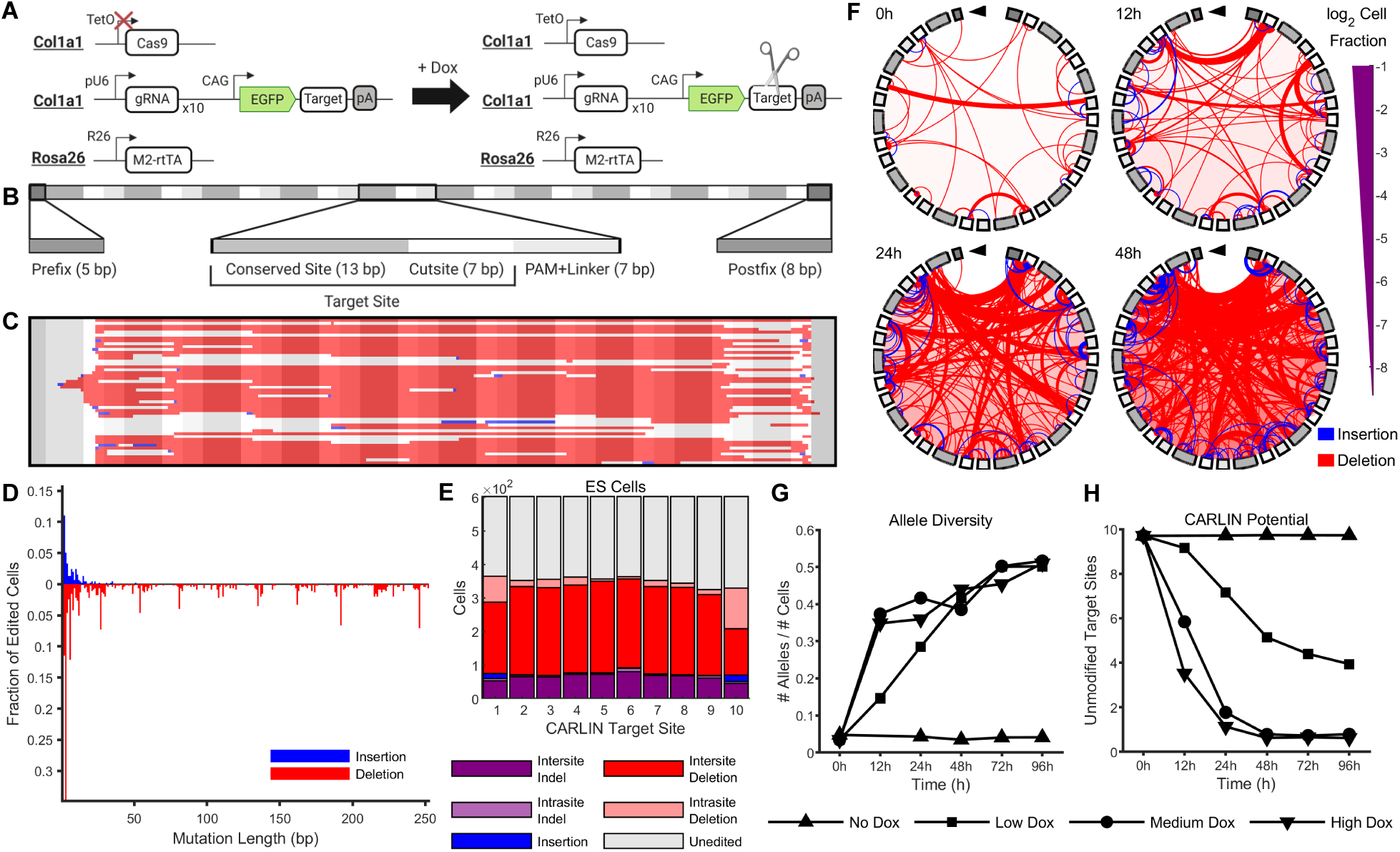
A high diversity of edits are generated by CARLIN in embryonic stem cells. **A.** Schematic of CARLIN system. Guides RNAs, target sites and inducible Cas9 components are contained within the *Col1a1* locus. The expression of each of the 10 gRNAs is driven by a separate U6 promoter (pU6). The CARLIN array sits in the 3’UTR of GFP and consists of 10 sites that perfectly match the gRNAs. The doxycycline (Dox) reverse tetracycline-controlled transactivator (rtTA) is contained within the *Rosa26* locus. **B.** For computational purposes, we consider the CARLIN array as a series of motifs. We divide each target site into a 13bp conserved site (that lies outside the expected range of Cas9 editing) and 7bp cutsite. Consecutive target sites are interleaved by a 3bp protospacer adjacent motif (PAM) and 4bp linker sequence. There is a 5bp prefix motif upstream of the first target site and an 8bp postfix motif downstream of the last target site. **C.** The 50 most common edited CARLIN alleles generated in CARLIN mouse embryonic stem (ES) cells following 96h induction with 0.04μg/mL Dox. Each row represents a different allele. Deletions are marked in red. Insertions are shown in blue with the left endpoint indicating the start of the insertion; the length of the strip matches the length of the insertion (except when occluded by a subsequent deletion). A grayscale mask as in (B) is overlaid to demarcate the CARLIN motifs. **D.** The fraction of edited ES cells, following 96h induction with 0.04μg/mL of Dox, in which insertions and deletions of various lengths are observed. **E.** Distribution of mutation types across different target sites in ES cells following 96h induction with 0.04μg/mL of Dox. **F.** Chord plots of CARLIN alleles before induction and at 12h, 24h, and 48h after induction with 0.04μg/mL of Dox. The shading of the iris (ccw. from top) corresponds to the shading of the motifs in Figure 1B (from left to right). The thickness of an interior line is proportional to the number of cells with that mutation. The endpoints of a red line indicate the starting and ending bps of a deletion. The upstream endpoint of a blue line indicates the insertion site, and the downstream endpoint is offset by an amount equal to the insertion length. **G.** Time-course of the total number of distinct alleles detected normalized by the total number of cells, and (**H**) average CARLIN potential (Methods) across cells in the absence of Dox and after induction with 0.04μg/mL (low), 0.2μg/mL (medium) and 1μg/mL (high) of Dox.

Taken together, doxycycline (Dox) induction drives Cas9 expression, which leads to double-strand DNA breaks in the target array. These breaks are repaired to result in a diverse range of altered DNA sequences (referred to as CARLIN alleles) that are expressed and stably inherited (Figure 1A). To analyze the CARLIN alleles from sequencing of the target array, we developed a novel bioinformatic pipeline that accounts for the location at which the Cas9-dependent alterations are expected (Methods; Figure 1B; Supplementary Figure 2A-C). This code is available at https://gitlab.com/hormozlab/carlin.

To test the ability of our system to generate inducible and detectable CARLIN alleles at the DNA level, we characterized the CARLIN edits present in CARLIN ES cells following Dox treatment (Methods). While we observed little background editing in the absence of Dox (Supplementary Figure 1C,F), a diverse set of repair outcomes was generated following Dox exposure (Figure 1C,D) as determined by high-throughput sequencing. These edits included deletions spanning 1-252 bps, the most common of which spanned multiples of 27 bps (the length of a target site and adjoining PAM+linker sequence), and insertions of up to 51 base pairs in length (Figure 1D). The edits occurred throughout the array, with different target sites displaying slightly different indel preferences (Figure 1E). In initial experiments, we observed 301 distinct alleles in 453 cells with edited alleles, 219 of which were only observed in one cell, indicating that CARLIN can generate highly diverse repair outcomes following Cas9 activity. To partially validate the results, we verified that the distribution of allele lengths produced by the bioinformatics pipeline was consistent with the distribution produced by fragment analysis (Supplementary Figure 2D), suggesting that the results are not distorted by library preparation, sequencing, or analysis steps.

We also investigated how CARLIN editing is influenced by the duration and magnitude of Cas9 expression. We exposed ES cells to low, medium and high dosages of Dox (0.04, 0.20 and 1.00μg/mL respectively), and performed bulk DNA sequencing prior to induction and at a series of timepoints up to 96h. As expected, both the fraction of cells with edited CARLIN sequences and the diversity of CARLIN alleles increases with both length and dose of induction (Figure 1F,G, Supplementary Figure 1C-F). This analysis also reveals that the nature of CARLIN edits can act as a readout of induction duration and strength. Specifically, we observe a decrease in the number of unmodified target sites (calculated as CARLIN potential, Methods) and an increase in the average length of deletions with increasing time and concentration of doxycycline (Figure 1H, Supplementary Figure 1F). Together, these data demonstrate that CARLIN is edited in an inducible way with the extent of editing dependent both on the duration and magnitude of the induction, indicating that the system can be used as a heritable molecular recorder.

### Sequential CARLIN induction permits lineage reconstruction

Having shown that we could regulate the extent and nature of editing, we next tested whether we could accrue sequential edits on the same CARLIN array. CARLIN ES cells were exposed to one, two or three 6h pulses of Dox (0.04μg/mL) interspersed by 24h in fresh media. Indeed, we observed an increase in the fraction of cells with edited CARLIN alleles, the number of mutations accrued in each allele, and the diversity of CARLIN alleles over the three pulses (Figure 2A; Supplementary Figure 1G). This finding indicates that sequential pulses of Dox can incorporate additional information and can potentially be used to build multi-level, hierarchical histories for lineage reconstruction. To test this last hypothesis, we exposed ES cells to one pulse of Dox, picked 8 ES cell clones for outgrowth, and exposed them to a second pulse of Dox (Figure 2B). Sanger sequencing of 8 ES cell clones after the first timepoint allowed us to establish a ‘ground truth’ of edits generated after the initial pulse (Figure 2C). We devised a basic lineage tree reconstruction algorithm that accounts for the expected CARLIN mutation patterns and assumes that the internal nodes are restricted to the observed alleles (Figure 2D; Methods). Applied to these data, we achieved a false positive rate of 0.6% (the fraction of cells in which the most recent ancestor of an allele is a clone other than where the allele came from) and a false negative rate of 18% (the fraction of cells in which none of the 8 selected clones is found as an ancestor of an allele). Therefore, sequential Dox pulses allow multiple stages of lineage reconstruction.

**Figure 2:**
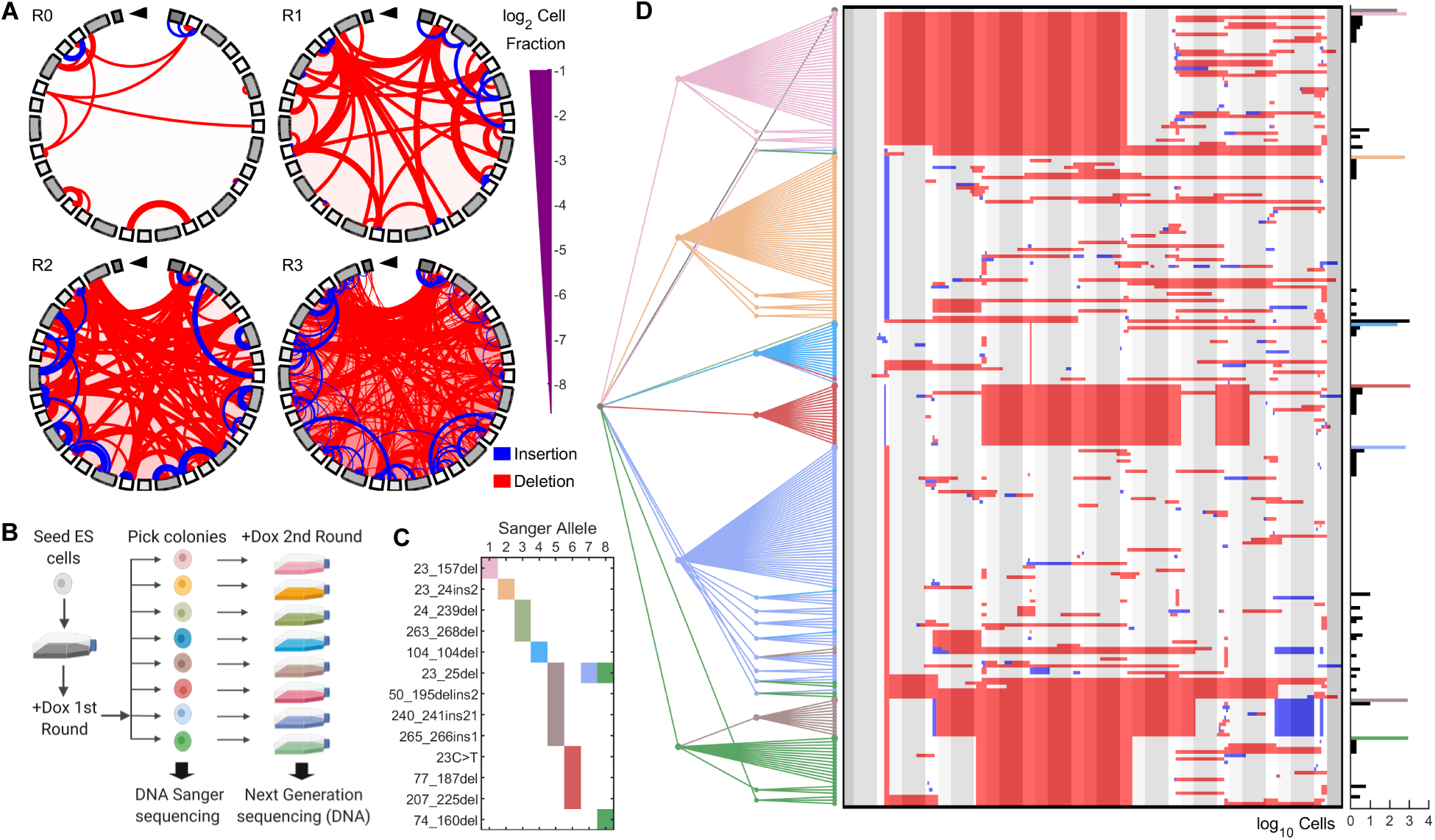
Multiple pulses of doxycycline can consecutively label lineages and enable phylogenetic tree reconstruction in embryonic stem cells. **A.** Chord plots of CARLIN alleles in the absence of doxycycline (Dox) and after one, two or three 6h pulses of Dox (R0-3, respectively). Color scheme as in Figure 1F. **B.** Following one 6h round of Dox induction, cells were seeded at single-cell density and 8 colonies were picked for further outgrowth and Sanger sequencing. Following a second round of Dox, DNA from cells was collected and sequenced by Next Generation Sequencing. **C.** Mutations called in each of the 8 colonies from the CARLIN pipeline applied to the Sanger sequences. Colonies are colored according to the schematic in (B). Colonies 5, 7 and 8 share a common mutation. **D.** (Left panel) The consensus tree, accounting for 95% of cells, obtained from 10,000 lineage reconstruction simulations applied to alleles pooled from all libraries (Methods, Supplementary Figure 4A). The color of a node and its branch to a parent corresponds to the NGS library in which the allele was observed. Leaves that connect to internal nodes of a different color correspond to false positives. (Centre panel) Sequence of each CARLIN allele visualized as in Figure 1C. (Right panel) Histogram of the number of cells in which each allele was detected. Colored bars correspond to NGS sequences which match a Sanger sequence.

### CARLIN generates a high diversity of barcodes in vivo

We next generated mice carrying the CARLIN transgenes and assessed allele generation following Dox induction in adults. Because dose and timing of Dox concentration is critical to induce editing in a large fraction of cells, we tested multiple dosing regimes of Dox (not shown) and selected a protocol in which CARLIN mice were exposed to Dox for 7 days (Methods). Following this protocol, we harvested RNA from multiple tissues from CARLIN mice for bulk sequencing of the CARLIN array (Figure 3A). We observed that the fraction of CARLIN transcripts that were edited ranged from 31% to 88% across all tissues analyzed, with the exception of the brain, that is inaccessible to Dox, and the heart and skeletal muscle, in which expression from the *Col1a1*/*Rosa26* loci has been previously shown to be low (Beard et al., 2006; Figure 3B, Supplementary Figure 3A). Importantly, background editing in the absence of Dox is negligible (averaging 1%) across all tissues of an uninduced 8-week old mouse (Figure 3B). Therefore, CARLIN represents a useful model to barcode adult tissues systematically.

**Figure 3:**
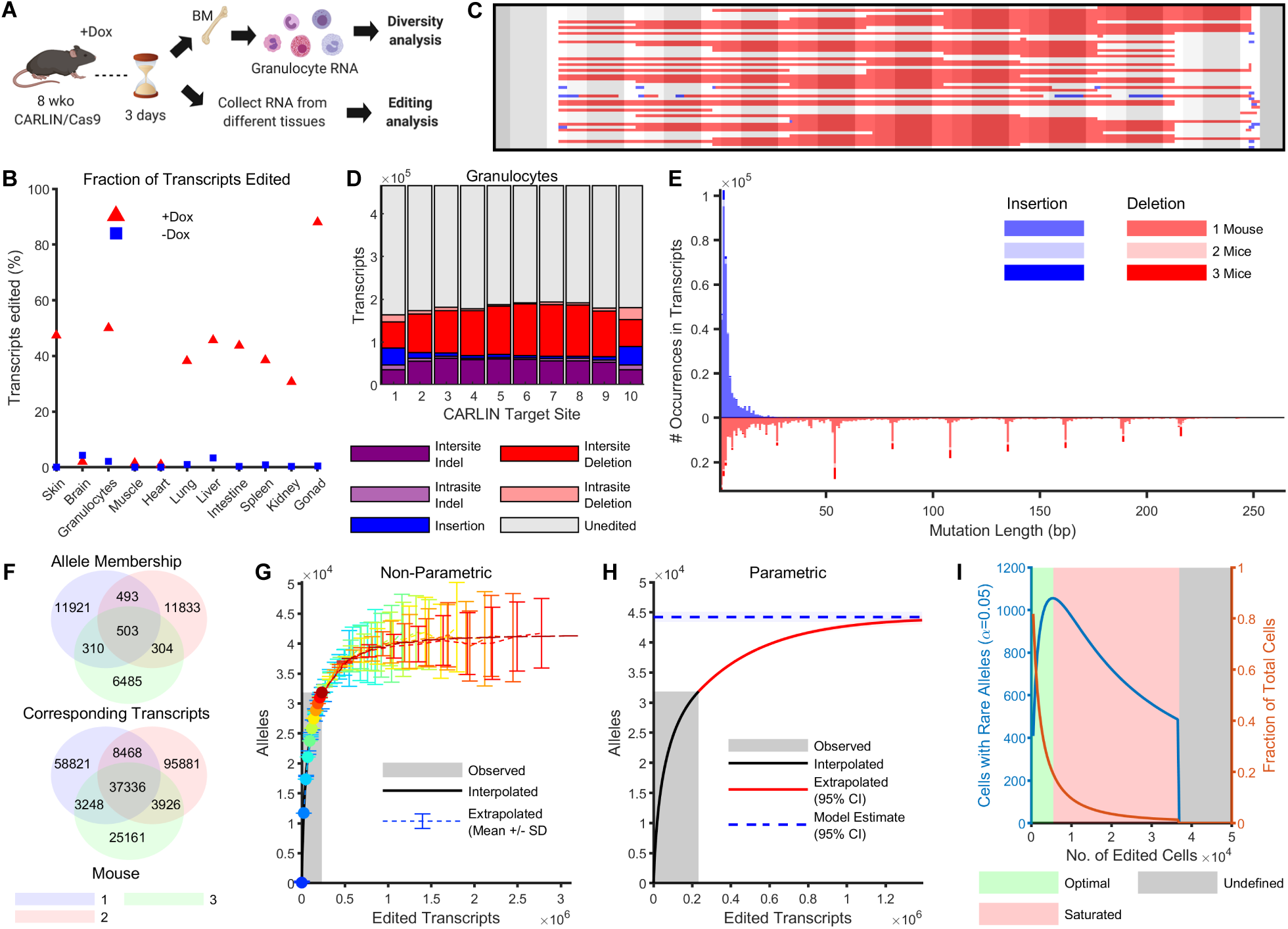
Inducible CARLIN editing *in vivo*. **A.** Eight week-old mice were induced with doxycycline (Dox) for one week. RNA from granulocytes and other tissues were collected following 3 days chase. **B.** Fraction of transcripts edited across tissues in the presence and absence of Dox. **C.** The 50 most common edited CARLIN alleles observed in granulocytes, visualized as in Figure 1C. **D.** Distribution of mutation types across different target sites in granulocytes comprising the allele bank (Methods). **E.** Histogram of insertion and deletion lengths found in the allele bank shaded according to presence across mice. **F.** Venn diagram showing number of edited alleles (and the corresponding number of edited transcripts) in the bank shared across the three induced mice. **G.** Non-parametric and (**H**) parametric extrapolation of the total allele diversity achievable by the CARLIN system as a function of the number of edited transcripts observed. The system is estimated to saturate at an allele diversity of 44,000 ± 400. The area shaded in grey indicates the number of observed transcripts used to construct the bank. **I.** Number of cells expected to harbor rare alleles (that are unlikely to occur independently in multiple cells) as a function of the number of cells edited. When the number of cells is small with respect to the CARLIN diversity (shaded in green), many cells harbor rare alleles. As the number of edited cells increases (shaded in red), the probability that a given allele marks only one cell decreases (orange curve), so that the number of cells that are uniquely marked with a CARLIN allele decreases (blue curve). In the regime shaded in grey, no cell can confidently be said to be uniquely marked by an allele (Methods).

We next assessed the full extent of the barcode diversity that could be generated *in vivo*. For this we compared CARLIN edits observed in large numbers of bone marrow granulocytes across three induced and two uninduced CARLIN mice following 1 week of Dox induction. Similar to our *in vitro* analysis, we detected a high diversity of edits in the induced mice generated through deletions and insertions across the length of the array (Figure 3C,D,E). Across the induced mice we observed the fraction of edited CARLIN transcripts ranging 29%-63%, compared to an average of 2.1% editing in the two uninduced mice (Supplementary Figure 3C,D). The editing in the uninduced mice is largely attributable to a low level of background Cas9 activity rather than resulting from errors introduced during library preparation, since editing in the absence of Cas9 was 0.3% (Supplementary Figure 3E). On average, 88% of the edited alleles (6485-11921) found in each mouse were not observed in other mice (Figure 3F), indicating that the majority of edits represent unique repair outcomes. However, it also indicated that a small percentage (∼12%) of alleles were generated at a higher frequency due to common indel mutation events (such as deletions spanning multiples of 27bps noted earlier) that independently generate the same allele sequence in different cells (Figure 3E). We pooled the edited alleles from across all induced mice together to form an allele bank, consisting of a total of ∼32,000 distinct edited alleles over ∼233,000 edited transcripts.

We used the allele bank to computationally estimate the total number of distinct alleles that CARLIN could generate (i.e. the maximum barcode diversity) and the expected occurrence frequency of these alleles. High diversity corresponds to many alleles that occur at equal frequencies. Conversely, low diversity corresponds to few dominant alleles that occur at high frequencies. By extrapolating the frequency distribution of alleles in the bank, we estimate that CARLIN is able to generate up to 44,000 ± 400 distinct alleles (Figure 3G,H; Methods), consistent with a high diversity system. Additionally, we used the bank to discriminate between rare alleles that occur at low frequencies and commonly-occurring alleles. To do so, we used the allele bank to estimate the probability that a CARLIN allele is unique for a given number of observed cells, obtaining a p-value of significance (Figure 3I; Methods). This discrimination is critical for any experiment to ensure that an allele detected in many cells is due to the shared lineage history of those cells, as opposed to independent CARLIN editing events that coincidentally produced the same allele. Critically, these statistical measures can be adjusted to account for other experimental parameters (such as number of cells in the system, number of detected CARLIN alleles, etc.) and may be applied to other CARLIN experiments.

Finally, we also investigated edited alleles generated in granulocytes of iCARLIN mice, in which the expression of the sgRNAs, as well as Cas9, is driven by a tetO promoter (Supplementary Figure 1B). Similar to the constitutive guide CARLIN system, we observed a high diversity of edits (Supplementary Figure 3F), indicating that this system may be used as well to label cells with even tighter inducibility and potentially shorter labelling windows.

### Lineage reconstruction in vivo

To investigate whether multiple rounds of CARLIN labelling could be used to gain insight into cellular phylogeny *in vivo* as done *in vitro* (Figure 2D), we set up timed pregnancies and delivered three pulses of Dox to pregnant dams at E6.5, E9.5 and E13.5 (Figure 4A). When the 3x labelled CARLIN embryos reached 8 weeks of age, we collected RNA from the skin, heart, liver, intestine and colon, and also separately sampled the left and right brain, muscle, lung and bone marrow HSCs, MPPs, granulocytes, and B-cells. We employed the same tree reconstruction algorithm developed for analyzing the *in vitro* experiment with ES cells (Figure 2D). However, as a pre-processing step we retained all alleles whose observed frequency was significantly higher than their frequency in the bank (using a FDR=0.05 on their frequency p-values – see Methods; Figure 4B). Only a small fraction of CARLIN transcripts were discarded based on this filtering step across all tissues (Figure 4C). We computed a consensus lineage tree, by simulating 10,000 stochastic reconstructions (Methods), which allowed us to visualize a hierarchy of clades across multiple tissues (Figure 4D-G, Methods). Based on this lineage tree, we computed a pairwise similarity matrix of the tissues and observed that contra-lateral tissues were closely related, as were multiple cell types of hematopoietic origin, and tissues of endodermal origin (Figure 4H), which is consistent with known lineage relationships. We also observed a low level of allele sharing across other tissues, some of which were derived from the same embryonic germ layer (Figure 4F,H). This indicates limited lineage mixing between these tissues and suggests that they began to develop independently prior to the stages of induction used in our analysis. Taken together, CARLIN can be useful for multi-level tissue reconstruction *in vivo*.

**Figure 4:**
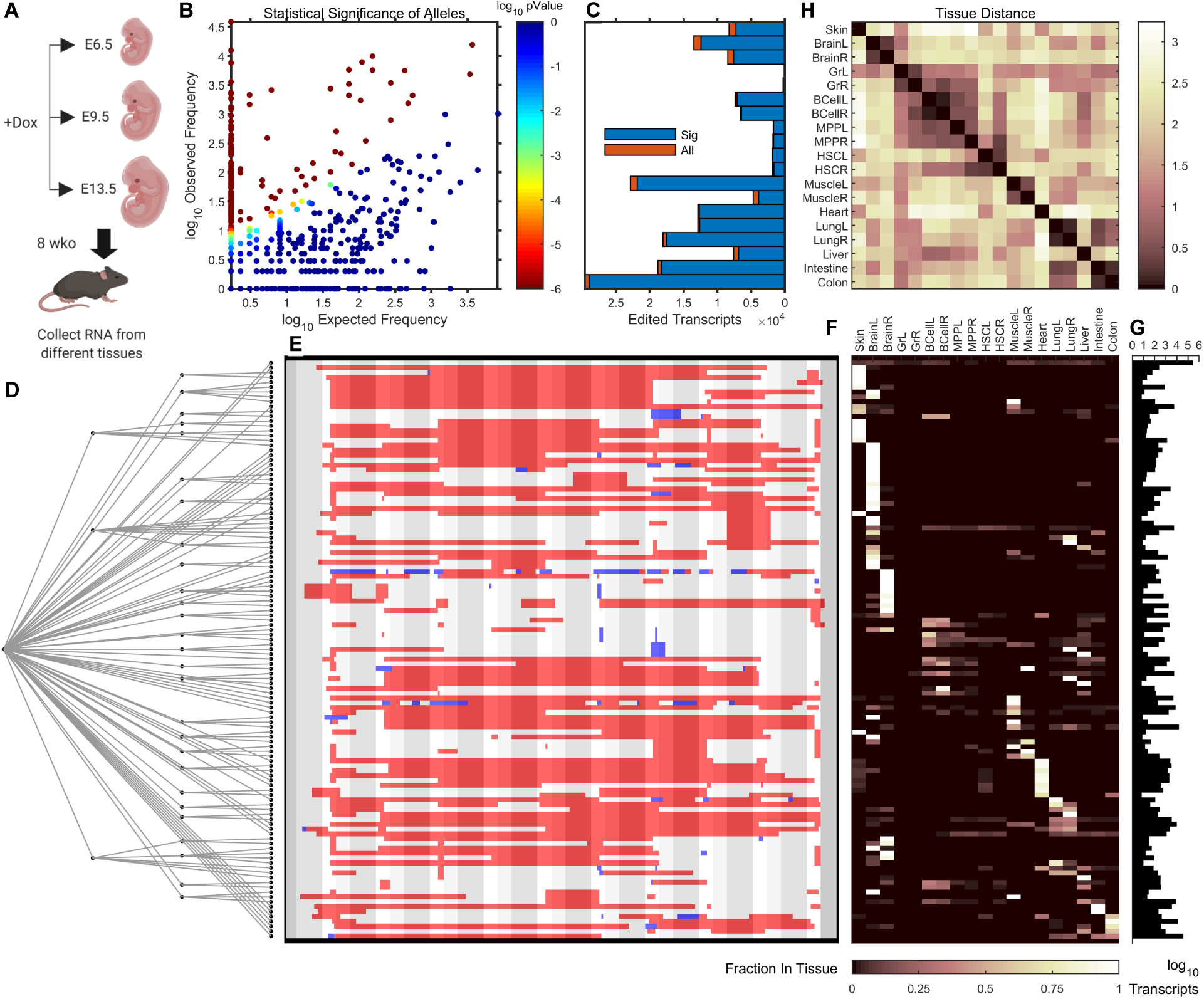
Lineage reconstruction *in vivo* through multiple pulses of doxycycline. **A.** Pregnant dams were induced with doxycycline at E6.5, E9.5 and E13.5. At 8 weeks, RNA from different tissues was collected and sequenced by Next Generation Sequencing. **B.** Scatter plot of observed allele frequencies vs. expected frequencies obtained by querying the bank. Alleles whose statistical significance did not survive a FDR of 0.05 were discarded (Methods). **C.** Number of edited transcripts found in different tissues after running the CARLIN pipeline (All), and after screening for significant alleles (Sig) as described in (B). **D.** The consensus tree which accounts for 95% of edited transcripts, obtained from 10,000 simulations, using the same algorithm as in Figure 2D (Supplementary Figure 4B; Methods). **E.** Allele sequences called from NGS corresponding to the leaf nodes, visualized as in Figure 1C. **F.** Distribution of number of transcripts corresponding to each allele across tissues (row normalized to 1). **G.** Histogram of total transcript counts across all tissues for each allele. **H.** Pairwise similarity matrix of tissues computed across alleles of the consensus tree (see Methods).

### Simultaneous detection of CARLIN barcodes and whole-transcriptomes from single cells

We next set out to develop a platform to detect CARLIN alleles using single-cell sequencing technologies. Our pipeline for this analysis involves: (i) exposure of mice to Dox, (ii) encapsulation of single cells from the cellular population of interest into droplets containing barcoded polyT-coated beads, (iii) amplification of whole cellular transcriptome, (iv) targeted amplification of the CARLIN array, and (v) sequencing using Next Generation Sequencing (Figure 5A). After optimization, we were able to detect CARLIN in 32-63% of cells in which we could also measure a full transcriptional profile (for the criteria used to select single cells, see Supplementary Table 3 and Methods). To check for reproducibility of our protocol, we prepared two CARLIN amplicon libraries independently starting from the same single-cell transcriptome library. We observed that 89% of the cell barcodes were shared across the two libraries. We also verified that the same CARLIN alleles occurred across the two samples with consistent frequencies (Supplementary Figure 6A, Methods).

**Figure 5:**
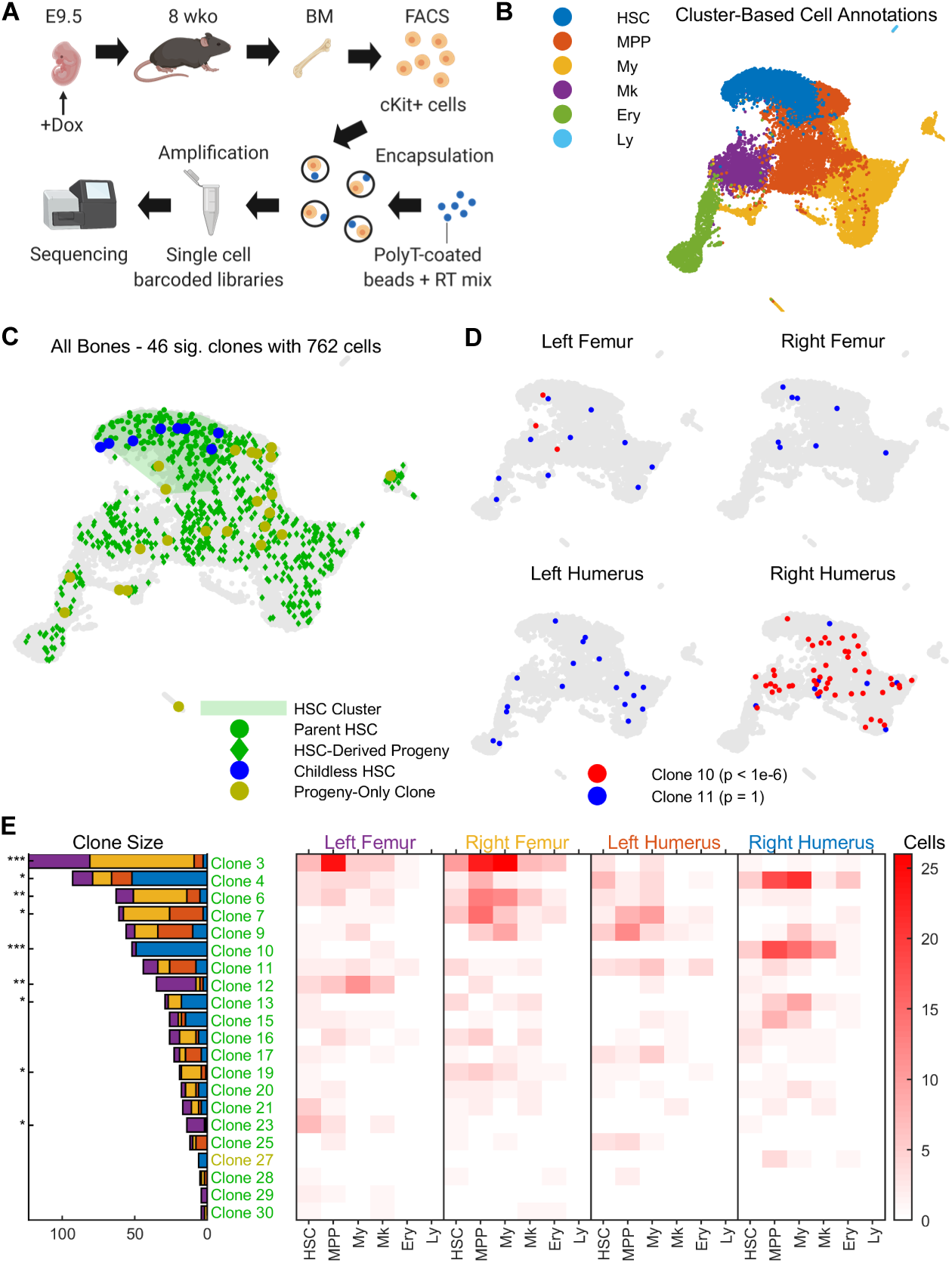
Clonal tracing of blood progenitors to adulthood. **A.** CARLIN mice were labelled at E9.5. At 8 weeks, bone marrow cells were collected, sorted, and encapsulated for single-cell RNA sequencing. **B.** UMAP representation of pooled transcriptome data from the bone marrow of 4 separate bones. See Supplementary Figure 5D,E for a breakdown of clusters and markers used for annotation. HSC, hematopoietic stem cell; MPP, multipotent progenitor cell; My, myeloid progenitor cells; Ery, erythrocyte; Ly, lymphoid cell. **C.** Statistically significant CARLIN alleles (FDR = 0.05; Methods) across all bones combined, overlaid onto the UMAP plot from (B). The green shaded area corresponds to the HSC cluster in the transcriptome, shown in (B). We are able to directly map the ancestry between differentiated cells (green diamonds) and HSCs (green circles) which share the same set of alleles. HSCs without children are shown in blue, and differentiated cells that do not share their allele with HSCs are shown in yellow. **D.** CARLIN clones overlaid onto the transcriptome of individual bones; a non-biased clone (blue) and a biased clone (red) is shown with the Bonferroni-corrected p-values for bone bias (Methods). **E.** (Left) Bar graph indicating the prevalence of each statistically significant allele across the 4 bones, with the Bonferroni-corrected p-value for bone bias marked as *p<0.05; ** p<10^−3^; ***p<10^−6^. (Right) Heatmap indicating occurrence frequency of alleles across bones and cell types. Alleles found in fewer than 4 cells are not displayed. The clone labels follow the color scheme in (C).

As a proof-of-principle experiment, we used CARLIN to characterize clonal properties of hematopoietic development. Here, we barcoded HSC precursors during embryogenesis and characterized their clonal lineages in adulthood. In the mouse, definitive blood progenitors arise at embryonic day (E) 10.5 with the formation of Runx1-expressing clusters within the main arteries of the embryo (Dzierzak and Bigas, 2018). From E11 onwards, these progenitors migrate to the fetal liver where they undergo extensive expansion before colonizing the bone marrow at around the time of birth. Although several studies have investigated the process through which the progenitors are formed, the dynamics of HSC expansion and migration to the bone marrow are still poorly resolved. In particular, it is unclear whether HSCs derived from the same developmental precursor clone already exhibit intrinsic functional biases.

We applied a single pulse of Dox at E9.5 to label the earliest emerging definitive blood progenitors (Figure 5A). Accounting for delays in Dox response and Cas9-protein stability (Alemany et al., 2018; Traykova-Brauch et al., 2008), this represents actual labelling times of approximately E10-E12.5. Once the labelled animals reached 8-weeks of age, we sorted a combination of Kit+ progenitors, including HSCs, multipotent progenitors (MPPs) and lineage-restricted progenitor cells (Supplementary Figure 5A) from four separate bones (right and left humerus and femur) and encapsulated the cells from each bone into separate single-cell libraries. We combined the 3755-5261 cells per bone that passed quality control cutoffs (Supplementary Table 3, Methods) into one dataset encompassing 19,056 cells (Supplementary Figure 5C). Unsupervised hierarchical clustering resulted in 28 distinct clusters that we annotated as HSC, MPP, myeloid, megakaryocyte, erythroid and lymphoid using previously described markers (Figure 5B, Supplementary Figure 5D,E). We considered cells belonging to the HSC-like cluster to be HSCs, and cells belonging to other clusters to be non-HSCs for all subsequent analysis. Finally, we visualized the single-cell gene expression profiles using uniform manifold approximation and projection (UMAP) plots, overlaid with the detected CARLIN alleles (Figure 5C).

From a total of 60 clones, each marked with a different CARLIN allele, our high-stringency analysis determined 46 (20-29 in each bone) to be significant (assessed using their clonal p-value at a FDR=0.05, see Methods; Supplementary Table 3). We restricted all further analysis to these significant clones. The sizes of these clones ranged from 1 to 123 cells comprising numerous cell types across the hematopoietic hierarchy. We initially assessed the extent to which clones that contained HSCs (HSC-rooted clones) also contained hematopoietic progeny (non-HSCs). Previous studies suggest that hematopoiesis is driven by HSCs that are progeny of definitive embryonic precursors (Dzierzak and Bigas, 2018). Indeed, we find that 23 out of 30 clones containing an HSC have detectable hematopoietic progeny (we refer to these HSCs as parent HSCs). Such HSC-rooted hematopoietic progeny make up most of the cellular composition in the analyzed bone marrow samples, i.e. 96% of non-HSCs displaying a significant CARLIN allele (p < 10^−6^; Figure 5C; Methods). Interestingly, there are 7 HSC clones for which we cannot detect hematopoietic progeny (Figure 5C). These ‘childless’ HSCs could potentially represent a novel HSC type that remains dormant throughout development and early adulthood. We also observed that the distribution of HSC-rooted clone sizes was significantly non-uniform (p<10^−6^; Figure 5E; Methods). Therefore, our data reveal highly heterogeneous parental outcome of embryonic-derived HSCs.

We next separated out the transcriptional and lineage profiles of cells across the four bones (Figure 5D). With this analysis we could assess both the presence and behavior of HSCs across multiple bone marrow compartments. We observed 13 of the 46 significant clones in all bones, accounting for 46% of cells displaying an edited CARLIN allele (Figure 5E). Notably, across all clones we observed that many of the largest clones were not uniformly distributed across bones, but were more likely to be found in a subset of the bones analyzed (Methods). For example, clone #10 appeared in 49 cells of the right humerus but appeared in only 3 cells of the left femur and was completely absent in the other analyzed bones (p < 10^−6^; Figure 5D,E). Similarly, clone #3 appeared in 42 and 72 cells of the left and right femur, respectively, but appeared in only 6 and 3 cells of the left and right humerus (p < 10^−6^; Figure 5E). No clone had a statistically significant fate bias, as judged by its occurrence among cell types as defined in Figure 5B, either within bones or pooled across all bones (Methods). Assuming equal expansion in the fetal liver, our data suggest that the expansion potential of fetal liver-derived HSCs might not be pre-determined but that it might be conferred by the niche into which they home. It is also possible that fetal liver-derived HSCs exhibit biases in colonization of different bones.

### Clonal bottlenecks during hematopoietic regeneration

Next, we applied CARLIN to investigate the clonal dynamics of adult hematopoiesis following perturbation. Decades of work have established that following acute myeloablation, most HSCs exit their quiescent state and undergo cell division (Harrison and Lerner, 1991; Wilson et al., 2008). It has been assumed that these divisions are asymmetric cell divisions and correspond to HSC activation, implying that most HSCs participate in regeneration. However, this process has never been studied at a clonal level and the extent to which each individual HSC participates in regeneration is unclear.

To measure how much individual HSCs contribute to regeneration, we studied the HSC response to 5-fluorouracil (5-FU), a widely used model of myeloablation in the mouse. 5-FU induces proliferation of most HSCs within 4 days (Harrison and Lerner, 1991; Wilson et al., 2008) and by 10 days post 5-FU most cellularity is recovered in the bone marrow (Harrison and Lerner, 1991). We induced CARLIN labelling in 8-week old mice before administering one dose of 5-FU via intraperitoneal injection (Figure 6A). Following 10 days recovery, we sorted the cKit+ population from the marrow of single bones (Supplementary Figure 5A,B) and generated single-cell RNA libraries. Across three independent experiments in control and 5-FU-treated groups, we detected between 4073-6025 cells with high resolution whole transcriptome information (Supplementary Table 3) that we compiled into one dataset following batch correction (Supplementary Figure 6B). Unsupervised hierarchical clustering resulted in 21 distinct clusters to which we assigned coarse-grain annotations as before (Figure 6B, Supplementary Figure 6C,D).

**Figure 6:**
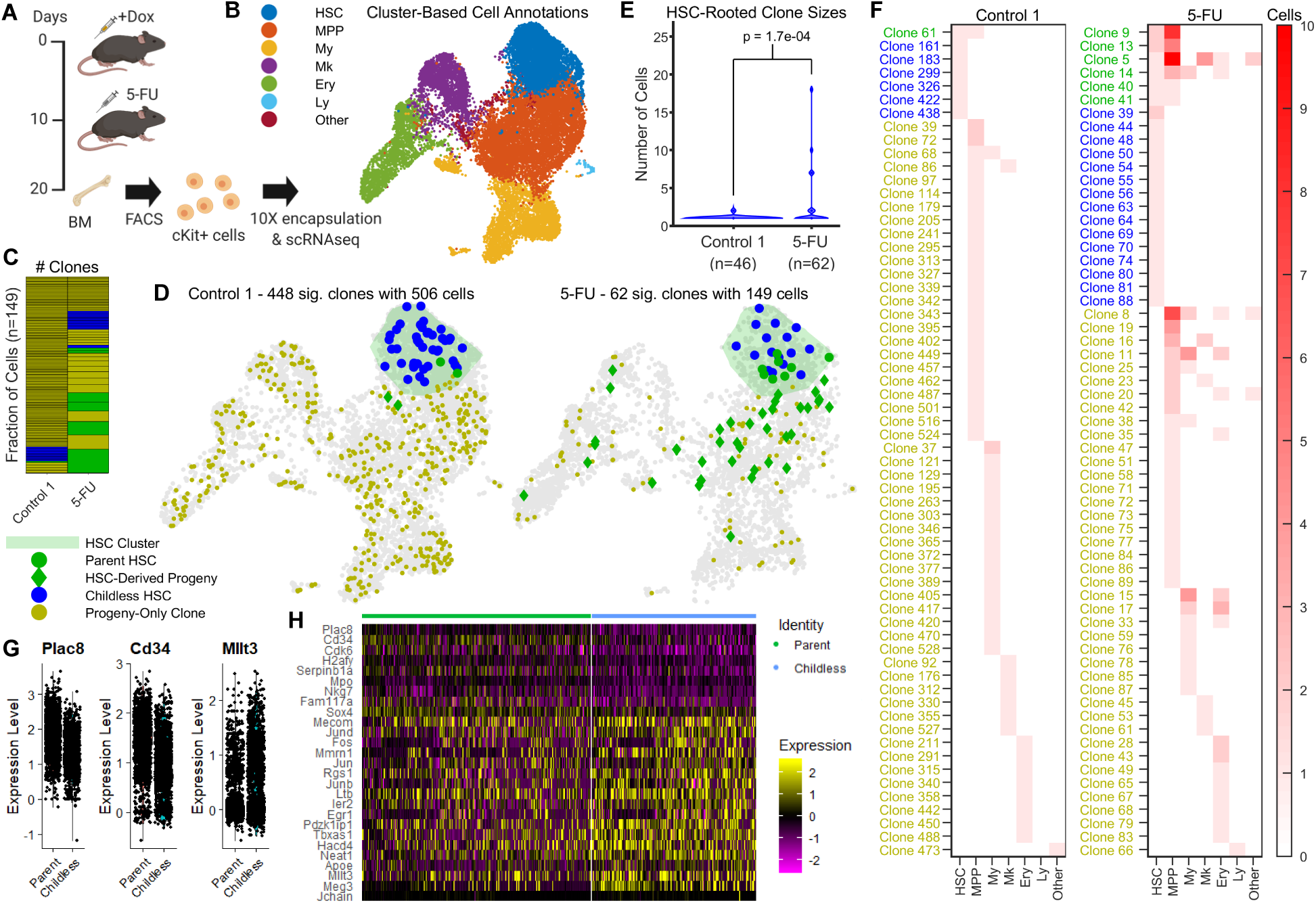
Clonal dynamics of adult hematopoiesis following perturbation. **A.** 8 week old CARLIN mice were labelled with doxycycline and injected with 5-FU after 10 days. After another 10 days, bone marrow cells were sorted and encapsulated for single-cell RNA sequencing. **B.** UMAP representation of pooled transcriptome data from control and 5-FU treated mice. See Supplementary Figure 6C,D for breakdown of clusters and markers used for annotation. Cluster labels as in Figure 5B. **C.** Number of statistically significant clones in the control and 5-FU treated mouse (FDR=0.05; Methods) after downsampling the control mouse to have the same number of cells marked by statistically significant alleles as the mouse treated with 5-FU. The control mouse has many small clones. The colors correspond to the legend for (D) below with blue clones containing only HSCs, yellow clones containing only non-HSCs, and green clones containing both. **D.** Statistically significant CARLIN alleles (as defined in C) overlaid onto the transcriptome indicating childless HSCs (blue), parent HSCs (green circles), non-HSC cells in an HSC-rooted clone (green diamonds) and non-HSC cells not in an HSC-rooted clone (yellow). The green shaded area corresponds to the HSC cluster in the transcriptome shown in (B). **E.** Violin plot showing the distribution of the number of cells in HSC-rooted clones in a control and 5-FU treated mouse (the green and blue markers in D). The total number of cells in HSC-rooted clones is shown in brackets under the sample label. **F.** Heatmap indicating occurrence frequency of statistically significant alleles (as defined in C) across different cell types in 5-FU and control animals. The clone labels are colored according to the scheme in (D). For the control, the number of clones have been downsampled to equal the number of clones in the 5-FU sample. **G.** Violin plots of selected differentially expressed genes between the parent and childless HSC cluster (as defined in Supplementary Figure 6E). **H.** Heatmap of differentially expressed genes between the parent and childless HSC cluster (as defined in Supplementary Figure 6E).

As with the previous experiment, we restrict our attention to clones corresponding to significant CARLIN alleles (assessed using their clonal p-value at a FDR=0.05, see Methods). We detected important differences in the clonal composition of control versus 5-FU treated bone marrow. First, we observed a significant reduction in the number of clones detected in the 5-FU-treated marrow (p<10^−6^; Figure 6C; Methods), which likely reflects the massive cellular and clonal loss after injury. Additionally, we used CARLIN to analyze the clonal contribution of HSCs to hematopoietic production. In the absence of 5-FU, only 20 of 1330 clones across all samples, representing 58 of 1522 (4%) edited CARLIN cells, contained both hematopoietic progeny and HSCs (Figure 6C,D,F). This suggests minimal contribution of HSCs at steady state, at least over 20 days, consistent with other studies (Busch et al., 2015; Sun et al., 2014). In the presence of 5-FU, this landscape was significantly altered with 6 of 62 clones containing both hematopoietic progeny and HSCs (p=1.7×10^−6^; Methods), representing 31% of cells carrying an edited CARLIN allele (p<10^−6^; Figure 6D,F; Methods). Additionally, there was a significant increase in the average size of HSC-rooted clones (p=1.7×10^−4^; Figure 6E, Supplementary Table 4; Methods). Surprisingly however, the distribution of the sizes of the HSC-rooted clones was significantly non-uniform (p=1.6×10^−3^; Figure 6E; Methods), with 4 of 21 HSC clones making up 68% of all cells in the HSC-rooted clones (Figure 6E). These findings indicate that a small number of highly-active HSCs are responsible for the replenishment of the blood system following cytotoxic injury. Therefore, our results indicate a clonal bottleneck during regeneration where only a handful of HSC clones can generate productive flow into the MPP and downstream compartments.

### CARLIN allows the identification of gene signatures underlying functional heterogeneity

As highlighted above, current clonal tracing models (Sun et al, 2014; Pei et al, 2017) in the hematopoietic system are able to identify heterogeneity in function. However, these studies cannot provide any molecular insight into potential drivers of function in HSCs. We explored whether CARLIN could allow us to identify gene signatures specific to the ‘active’ HSC state. Initially, we performed differential gene expression analysis comparing the parent HSCs (n=35) to childless HSCs (n=242) across the control and 5-FU single-cell datasets (Supplementary Table 5). However, only one of these genes, *Plac8*, showed a statistically significant change at a log-fold change cutoff of 0.25, after applying a Bonferroni correction (average log fold change=0.44, p=2.2×10^−2^). To increase the number of cells used for the differential analysis, we took advantage of our observation that, as visualized using UMAP, the parent HSCs were not uniformly spread across the HSC Louvain cluster (Figure 6B,D). To delineate the parent HSC region in an unbiased way, we changed the Louvain clustering resolution parameter and obtained two sub-clusters within the original HSC cluster, one of which contained a significantly larger fraction of parent HSCs compared with the other (z=2.3, p=1.2×10^−2^; Supplementary Figure 6E; Methods). Differential gene expression analysis across these two sub-clusters revealed a higher number of significantly different genes, in addition to *Plac8* (Figure 6G,H, Supplementary Figure 6F,G, Supplementary Table 6). Some of these genes have known involvement in HSC quiescence/ activity (*Mllt3, Cd34, Pdzk1ip1*; Forsberg et al., 2010; Pina et al., 2008; Wilson et al., 2008), hematopoietic differentiation (*Mpo*) and cell proliferation (*Cdk6, Plac8;* Rogulski et al., 2005), as well as a number of genes with described but poorly-defined links to hematopoiesis (*Nkg7, Fam117a*; Wilson et al., 2015). Taken together, these data demonstrate that the combined analysis of lineage and gene expression profiles can in principle identify molecular profiles underlying heterogeneous HSC behavior *in vivo*, without a need for *a priori* known markers or cell sorting.

## Discussion

Here, we present CARLIN, a new resource for lineage tracing research that can be used to simultaneously interrogate lineage histories and gene expression information of single cells in the mouse in an unbiased, global manner. We have demonstrated that CARLIN mice can be used to generate up to 44,000 distinct CARLIN alleles (barcodes) *in vivo*, and that these alleles can be detected and read out using single-cell droplet sequencing alongside the transcriptome of individual cells. We also demonstrated that multiple pulses of labelling can be used to enhance our understanding of tissue phylogeny.

CARLIN has a number of unique advantages over existing mouse lines for *in vivo* lineage tracing. Unlike models that use Polylox (Pei et al, 2017) or Sleeping Beauty transposons (Sun et al, 2014), the barcodes generated by CARLIN are transcribed, enabling i) high-throughput readout of lineage histories in single cells and ii) simultaneous measurement of gene expression profiles in the same cells. Because of this, we can characterize the identity of the cells that have been traced using their gene expression profiles in a precise and unbiased fashion. In contrast, existing techniques sort cells into subpopulations based on known cell types prior to readout of lineage histories. This requires prior knowledge of the markers associated with each cell type of interest and existence of antibodies that can enrich for these subpopulations. Strategies that rely on cell type specific expression of fluorescent reporters require costly and time consuming genetic engineering. Even with available established cell sorting strategies, resulting cell purity is limited and cells can be lost during sorting. Critically, CARLIN can be used to read out the lineage histories of any cell type, in any tissue and organ, even in the absence of known cell surface markers for sorting. Therefore, CARLIN enables precise annotation of cell types whose lineage has been traced beyond what can be gleaned from cell surface markers alone. Complete gene expression profiles also provide information about mechanisms that drive cell behaviour. Finally, CARLIN can directly quantify clone sizes by counting the occurrence frequency of barcodes in individual cells. Existing methods rely on bulk sequencing and are therefore less accurate because of PCR amplification biases.

Innovations that have increased our ability to both modify and detect DNA sequences in single cells has led to sophisticated lineage tracing systems based on CRISPR-Cas9 gene editing, with recent implementations in mice (Chan et al., 2019; Kalhor et al., 2018). In these previously described models, constitutive Cas9 expression generates evolvable barcodes in hundreds of random genomic target sites. Our system offers a number of advantages over these previously described models. First, our system is inducible, allowing cells to be barcoded at precise timepoints. Second, all transgenic elements in CARLIN mice are contained in defined genomic loci, enabling straightforward crossing into alternative genetic backgrounds, and minimizing damage caused by continuous double-strand DNA breaks. Third, we have created a bank of alleles and statistical methods to create a confidence score for each allele, allowing us to quantify the statistical significance of alleles that are shared across multiple cells. Finally, CARLIN mice represent a stable and practical mouse line that can be utilized by others in the scientific community, avoiding the use of zygote microinjection or complex mouse crosses.

We have used CARLIN to shed light on two aspects of hematopoiesis. First, we applied our tool to track early blood progenitor clones to adulthood. A surprising observation was that the majority of the largest clones detected exhibited significant bias in their representation across the four separate bones analyzed. One possible explanation for this is heterogeneity in the niche environments resulting in different seeding success and subsequent expansion (Gao et al., 2018). Alternatively, our data could indicate that only a subset of HSCs in the fetal liver expand or seed the bone marrow (Ganuza et al., 2017). Finally, we also cannot rule out that migration of HSC clones during adulthood occurs between bones, contributing to a skewed distribution of clones (Wright et al., 2001).

Second, we used CARLIN to analyze the clonal dynamics of blood replenishment following chemotherapeutic lympho/myeloablation. Analysis of CARLIN alleles uncovered a reduced clonal diversity of the blood following 5-FU treatment. Furthermore, we observed that a small number of HSCs were responsible for replenishment of the blood. This finding is surprising given that previous reports indicated homogenous cycling within the HSC compartment following 5-FU treatment (Harrison and Lerner, 1991; Wilson et al., 2008). Taken together, it is possible that 5-FU treatment could initiate widespread cycling within the HSC niche with only a small number of clones continuing to cycle and contributing to downstream blood populations. Interestingly, similar dynamics were observed following transplantation into irradiated or cKit-depleted mice (Lu et al., 2019) where only a subset of HSCS replenished the blood, suggesting that skewed blood production from HSCs could be a generalized response to hematopoietic stress. Differential gene expression analysis comparing these ‘active’ HSCs with their ‘inactive’ counterparts revealed increased expression of cell cycle and cell differentiation genes among others; of particular interest is *Plac8* that has been previously implicated in proliferation (Rogulski et al., 2005), host defence (Ledford et al., 2007) and has reduced expression in aged HSCs (Mann et al., 2018). Our identification of a potential new candidate gene involved in the regulation of HSC quiescence/ activation highlights the value of using CARLIN to interrogate the molecular drivers underlying the heterogeneous clonal output of HSCs.

Our method can potentially be improved in several ways. First, while our diversity estimates have established the maximum diversity of the CARLIN system to be ∼44,000 alleles, which is sufficient for many applications, higher diversity may be desired when analyzing whole adult tissues. Diversity of the system could be increased through simple modifications such as use of homing guide RNAs (Kalhor et al., 2018) or combining the system with Cre-based tracing lines. Second, the current iteration of CARLIN can result in a limited fraction of cells edited (16-74% of cells have edited CARLIN alleles across our single-cell datasets). Editing efficiency could potentially be increased by optimizing the timing and/or dose of doxycycline. Third, a CARLIN capture rate of 32-63% (fraction of cells in which CARLIN was detected across our single-cell datasets) may be limited by low expression/stability of CARLIN RNA, transcriptional bursting from the promoter used, or errors in PCR/sequencing resulting in loss of reads *in silico*. Incorporating CARLIN into loci that are more highly expressed, using additional CARLIN arrays, or further optimizing the promoter or RNA stabilization sequences could potentially circumvent these deficiencies and increase the fraction of cells from which lineage histories can be extracted.

Finally, this work among others (Alemany et al., 2018; Chan et al., 2019; Frieda et al., 2017; Kalhor et al., 2018; Raj et al., 2018; Spanjaard et al., 2018) represents a proof-of-principle study for the robust recording of cellular information using genome editing. In principle, CARLIN can be extended so that Cas9 expression is controlled by environmentally-sensitive promoters rather than doxycycline. Such a system could record histories of specific stimuli such as pathogen exposure, nutrient intake and signaling activity, in addition to lineage.

## Acknowledgements

We thank members of the Camargo lab and Dr. Donna Neuberg for helpful discussions. We also thank Ronald Mathieu for help with FACS and flow cytometry. DS was funded in part by the Natural Sciences and Engineering Research Council of Canada (NSERC PGSD2-517131-2018). SB and FGO were funded by EMBO (ALTF 798-2018 and ALTF 655-2016, respectively). This study was supported by awards from the National Institute of Health (HL128850-01A1 and P01HL13147 to F.D.C). F.D.C. is a Leukemia and Lymphoma Society and a Howard Hughes Medical Institute Scholar. SHO acknowledges support from the NIDDK-supported Cooperative Centers of Excellence in Hematology (CCEH) at BCH (U54 DK110805). SH acknowledges support from NIH R00GM118910 and the Harvard University William F. Milton Fund.

