## Supplementary Materials for "An engineered CRISPR/Cas9 mouse line for simultaneous readout of lineage histories and gene expression profiles in single cells"

**SUPPLEMENTARY MATERIAL**

**Methods**

Experimental Methods

Quantification and Analysis

**Supplementary Figures**

Supplementary Figure 1: Additional quantification of editing in ES cells

Supplementary Figure 2: Comparison of CARLIN analysis pipeline with NW alignment and fragment analysis

Supplementary Figure 3: Mutations in different tissues, and additional details of the allele bank

Supplementary Figure 4: Additional details of lineage tree reconstruction simulations

Supplementary Figure 5: FACS plots and extended data for embryonic induction experiment

Supplementary Figure 6: Extended data for 5-FU experiment

**Supplementary Tables**

Supplementary Table 1: Related to CARLIN construct design (annotated 5' to 3')

Supplementary Table 2: Penalties ($P_B/P_E$) used in alignment algorithm (Supplementary Figure 2 & Methods)

Supplementary Table 3: Metrics for single cell data (Figures 5 & 6)

Supplementary Table 4: Statistical significance (p-value) of differences in average HSC-rooted clone size (Figure 6)

Supplementary Table 5: Differential gene expression between parent HSCs and childless HSCs (Figure 6)

Supplementary Table 6: Differential gene expression between parent and childless HSC cluster (Figure 6)

#### EXPERIMENTAL METHODS

##### Contact for Reagent and Resource Sharing

##### Experimental Model and Subject Details

8-week-old mice (*Mus musculus*) were used in all the experiments unless noted in the text or figure legends. Both male and female mice were used indistinctly as we have not observed any difference associated with sex in the biological processes studied; mice were randomly assigned to the different experimental groups in the experiments shown. CARLIN and Cas9 mice were derived from the KH2 mouse embryonic stem cell (ESC) line, with a mixed C57BL/6 x 129 genetic background. Experimental mice used in this study were from F2/F3 generations resulting from the breeding of F1-C57BL/6 x 129 with the TetO-Cas9 mice (also mixed C57BL/6 x 129 genetic background). All mice were maintained in standard conditions of housing and husbandry at Boston Children's Hospital, and all the procedures involving animals were approved by the Boston Children's Hospital Institutional Animal Care and Use Committee.

The CARLIN mouse embryonic stem cell (ESC) line used in some experiments was derived from a male embryo from the F3 generation. Karyotype analysis was performed on the cells to ensure proper genome stability. ESCs were maintained in KO-DMEM, supplemented with 15% ES-FBS, 10 ng/mL LIF and non-essential amino-acids, and grown over a mono-layer of mitomycin C-inactivated mouse embryonic fibroblasts (MEFs), for the time points indicated in the results and figure legend sections. All cultures were maintained in standard tissue culture conditions of 37°C and 5% $CO_2$.

##### Design and Assembly of CARLIN Array

The CARLIN reference sequence was designed as an array of ten sense-oriented CRISPR/Cas9 target sites, each with a length of 20 bp and separated from each other by a 3 bp protospacer adjacent motif (PAM) sequence and 4 bp linker. In order to design a mouse-optimized array containing 10 CRISPR/SpCas9 target sites, we started from the set of guides previously tested for the zebrafish GESTALT system development (McKenna et al., 2016; v6 and v7 arrays). From that set, we first excluded all target sites showing any homology with the mouse genome. Secondly, we used CRISPR design tool (crispr.mit.edu) to select guides with the strongest score factor (highest efficiency and lowest off-target) possible. Based on these criteria we preserved 6 guides from the GESTALT system and designed from the scratch the other 4 using the same criteria described. To test each guide, we cloned each one of the 10 sgRNAs individually under the control of the U6 promoter and transduced along with a plasmid containing the Carlin array into 293T cells. 48 hours after transduction we lysed cells and PCR amplified the array for fragment analysis. All guides selected for the final design showed a similar activity in this assay, assessing the further steps performed with the array.

For array synthesis, the final sequence was synthesized as a gBlock (IDT) and is provided in Supplementary Table 1. The CARLIN reference sequence was cloned into the 3'-UTR region of a GFP open reading frame in an intermediate cloning vector, upstream of the bGH-polyA sequence, and under the control of the CAG promoter to ensure robust expression.

**Design and Cloning of the CARLIN sgRNA Multiplexes**

10 sgRNAs perfectly matching the target sites of the CARLIN reference sequence were expressed as a multiplex, in which each sgRNA is driven by its own promoter. We assembled two different multiplexes, one with Dox-inducible sgRNAs (iCARLIN; see Supplementary Figure 1B), and one with constitutive sgRNAs (constitutive CARLIN, used for all data in the paper with the exception of Supplementary Figure 3F), using the same Golden Gate assembly strategy. In the inducible multiplex, we used the FgH1tUTG donor plasmid (Addgene plasmid #70183) (Aubrey et al., 2015) to clone each one of the sgRNAs using the BsmBI restriction sites, whereas in the constitutive multiplex, we used the pU6-(BbsI)_CBh-Cas9-T2A-mCherry plasmid (Addgene plasmid #64324) (Chu et al., 2015). Then, the 10 blocks containing the promoter and the sgRNA were PCR-amplified using specific primers with overhangs for the Golden Gate assembly method. The Golden Gate assembly protocol was adapted from the Addgene protocol (https://media.addgene.org/cms/files/GoldenGateTALAssembly2011.pdf) (Cermak et al., 2011). All primer sequences are provided in Supplementary Table 1.

**Targeting ESCs with the CARLIN Transgene**

Both the CARLIN reference sequence and the sgRNA multiplex were cloned adjacently into the pBS31 targeting vector (Beard et al., 2006) to target the ESCs in the *Col1a1* locus. For an efficient targeting of the transgene in this locus we used the KH2 ESC line, containing a donor FRT site in the *Col1a1* locus, as well as the M2-rtTA transgene introduced in the *Rosa26* locus to allow expression of Dox-inducible systems. Approximately $1.5 \times 10^7$ KH2 ESCs were electroporated using 50 μg of the pBS31-CARLIN vector and 25 μg of pCAGGS-FLPe-puro (Buchholz et al., 1998) at 240 V and 500 μF using a Gene PulserII (Bio- Rad, Hercules, CA). Hygromycin selection (140 μg/ml) was started 24h after electroporation. The genomic DNAs from the selected clones were screened by PCR using the *Col1a1* genotyping primers (listed in Supplementary Table 1). The same protocol was independently performed for the inducible and constitutive CARLIN systems.

**Mice Generation**

Two targeted ESCs clones were selected from each of the inducible and constitutive CARLIN systems and injected into BL/6 embryos to form mouse chimeras. At least two chimeras with >90% of chimerism from each ESC clone were used as founders of our experimental mouse cohort. The ESCs were injected at the Mouse Gene Manipulation Core at Boston Children's Hospital. To generate experimental mice, descendants from the F1 of the injected ESCs were bred with *Col1a1-TetO-SpCas9*, generated at Stuart H. Orkin's laboratory. Primers for genotyping the *Col1a1* and *Rosa26* loci are listed in Supplementary Table 1.

#### Animal Procedures

Unless noted elsewhere, Dox was administered to 8-week-old mice via drinking water (2 mg/mL supplemented with 10 mg/mL sucrose) and three intraperitoneal injections (50μg/g) every other day. For the embryonic labeling, Dox was administered to pregnant females via retro-orbital (RO) injection with 25 μg/g of a 10mg/mL solution of Dox at E9.5. 5-fluorouracil (5-FU) was intraperitoneally injected (150 mg/kg) into 8-week-old mice. To prepare 10 mL of the 15 mg/mL 5-FU injectable solution, the chemical was first suspended in 500 μL NaOH 1N and then dissolved in 9.5 mL of phosphate-buffered saline (PBS). For peripheral blood extraction (used to assess fraction of CARLIN sequences edited before performing the single-cell experiments), mice were anesthetized using isoflurane and 2-3 capillaries were collected from the RO sinus. For bone marrow preparation, mice were euthanized and bones from the four limbs were immediately dissected and flushed with 2% fetal bovine serum in PBS. Erythrocytes were removed by osmotic lysis before antibody staining and fluorescence-activated cell sorting.

#### Fluorescence-Activated Cell Sorting

Unless noted, lineage depletion was performed in the whole bone marrow samples using magnetic-assisted cell sorting (Miltenyi Biotec) using the biotin-conjugated lineage markers CD3e, CD19, Gr1, Mac1, and Ter119. Cell populations were purified using FACSAria (Becton Dickinson) and the flow cytometry data was analyzed using FlowJo (Tree Star). The following combinations of cell surface markers were used to define the analyzed populations: LT-HSC: $Lin^-Kit^+Sca1^+CD150^-CD48$-; MPP3/4: $Lin^-Kit^+Sca1^+CD150^-CD48^+$; ST-HSC: $Lin^-Kit^+Sca1^+CD150^-CD48^-$; MPP2: $Lin^-Kit^+Sca1^+CD150^+CD48^+$; MyP: $Lin^-Kit^+Sca1^-CD150^-CD41^-$; MkP: $Lin^-Kit^+Sca1^-CD150^+CD41^+$; granulocytes: $Ly6G^+Mac1^+B220^-Ter119^-$; monocytes: $Ly6G^-Mac1^+B220^-Ter119^-$; pro-pre-B cells: $Ly6G^-Mac1^-B220^+Ter119^-$; erythroblasts: $Ly6G^+Mac1^+B220^-Ter119^+CD71^+$. Representative examples of sorted populations can be found in Supplementary Figure 5A,B. The following antibodies were used in this study at a 1:100 dilution: biotin anti-mouse CD3e [145-2C11] (eBioscience), biotin anti-mouse CD19 [MB19-1] (eBioscience), biotin anti-mouse Gr1 [RB6-685] (eBioscience), biotin anti-mouse CD11b (Mac1) [M1/70] (eBioscience), biotin anti-mouse Ter119 [TER119] (eBioscience), anti-biotin microbeads (Miltenyi Biotec), APC anti-mouse CD117 (c-kit) [2B8] (BioLegend), PE-Cy7 anti-mouse Ly-6A/E (Sca-1) [D7] (eBioscience), APC-Cy7 anti-mouse CD48 [HM48-1] (BD Pharmingen), PE-Cy5 anti-mouse CD150 [TC15-12F12.2] (BioLegend), BV605 anti-mouse CD41 [MWReg30] (BioLegend), eFluor450 anti-streptavidin (eBioscience), APC/Cy7 anti-mouse/human CD45R/B220 [RA3-6B2] (BioLegend), PE anti-mouse Ly-6G [1A8] (BioLegend), APC-Cy7 ant-mouse CD11b (Mac1) [ICRF44] (BioLegend), eFluor 450 anti-mouse CD3 [17A2] (eBioscience), PE-Cyanine5 anti-mouse Ter-119 [TER-119] (eBioscience), BV510 anti-mouse CD71 [C2] (BD Pharmingen).

#### CARLIN Amplification and Bulk Sequencing Protocols

The sequences of all primers listed here are shown in Supplementary Table 1. For all applications, the PCR amplification of the CARLIN array was performed using primers flanking the 5' and 3' external regions of the array (primers CARLIN_fwd1, CARLIN_fwd2 and CARLIN_rev). Illumina adaptor regions, unique motif identifiers (UMI), biotin and fluorescent tags are added to these primers according to the experiment needs. For fragment analysis, the CARLIN_fwd2 was conjugated with 6-carboxyfluorescein (6-FAM) in the 5' position (FAM_CARLIN_fwd), and PCR products were separated by capillary electrophoresis.

For sequencing the CARLIN array from the genomic DNA (gDNA) of pooled cellular populations, up to 250 ng of input gDNA were used per library. We first performed a UMI-tagging reaction using the CARLIN_fwd2 primer attached to the Illumina sequencing adapter and a 10 bp region of fully degenerate DNA sequence (primer NGS_UIM_D_F). This reaction was performed as a single extension step, in which temperature ramped between annealing and extension for five cycles without a denaturalization step to prevent re-sampling of gDNA CARLIN sequences (McKenna et al., 2016). The DNA product was then purified using AMPure XP beads (Beckman Coulter) and the whole volume was loaded into a PCR reaction to amplify UMI-tagged CARLIN sequences (primers NGS2_F, NGS1_R; 35 cycles). Finally, an indexing PCR was performed (primers P5, NGS2R_I#; 10 cycles) before sequencing.

For preparing libraries from RNA of pooled cellular populations, up to 1μg of total RNA was retro-transcribed using a gene specific primer that contains a 10 bp region of fully degenerated sequence (RT_Bulk_CARLIN) and SuperScript III (Invitrogen). For all *in vitro* applications, this cDNA product was then purified using AMPure XP beads (1X; Beckman Coulter) and loaded into a PCR reaction for CARLIN amplification (primers NGS2_F, NGS1_Bulk_R; 35 cycles; Platinum Taq enzyme (Invitrogen)). Following AMPure XP bead purification (0.8X), the indexing PCR was performed (primers P5, NGS2R_I#; 10 cycles). For all *in vivo* applications, the protocol was optimized for a nested PCR approach. The purified cDNA product was loaded into a first PCR reaction (primer NGS1_F, NGS1_Bulk_R; 15 cycles; Q5 High Fidelity Polymerase (M0491, New England Biosciences)). Following AMPure XP beads purification (0.8X), half the product was loaded into a second PCR reaction (primer NGS2_F, NGS1_Bulk_R; 15 cycles; Q5 High Fidelity Polymerase). Finally, after AMPure XP bead purification (0.8X) the indexing PCR was performed (primers P5, NGS2R_I#; 9 cycles). The final indexed libraries were purified with AMPure XP bead purification (0.8X). Libraries were sequenced on Illumina MiSeq using paired-end 500 cycles v2 kits (Read 1: 250 cycles; Index Read: 6 cycles; Read 2: 250 cycles) with 10% PhiX sequencing control v3 (Illumina).

#### Single-Cell RNA Sequencing Protocols

10X Chromium single cell 3' (10X) protocols were performed following the manufacturer's instructions and step-by-step protocols can be found at the companies' websites. For the 10X Chromium Single Cell 3' libraries, whole-transcriptome libraries were prepared following the 10X v2 (Figure 5; https://bit.ly/2OUeaUj) and v3 protocol (Figure 6; https://bit.ly/2YP9LoI). Whole-transcriptome libraries were sequenced on Illumina NextSeq500 using paired-end 150 cycles v3 kits (Read 1: 28 cycles; Index Read 1 (i7): 8 cycles; Read 2: 91 cycles).

For targeted CARLIN amplification, 3-5 ng of the amplified cDNA (Step 2.3) was loaded into an initial PCR reaction (10X-CARLIN_1-bio, P5-PR1; 15 cycles). Following cleanup with 0.8X

SPRIselect (Beckman Coulter), biotin-tagged products were purified using Dynabeads kilobaseBINDER Kit (Invitrogen) following the manufacturer's instructions. In short, Dynabeads were incubated with the PCR product on a roller for 3h at room temperature before supernatant removal using magnet separation and washing. Half of the CARLIN-tagged Dynabeads were then loaded into a second PCR reaction (10X-CARLIN_2, P5-PR1; 15 cycles). Following Dynabead removal by magnet, libraries were purified with 0.8X SPRIselect and the indexing step was performed using one tenth of the PCR product (SI primer, Chromium i7 sample primer; 8 cycles). For all 10X PCR reactions, the polymerase supplied with the 10X v3 kit was used. 10X CARLIN single-cell libraries were sequenced on Illumina MiSeq using paired-end 500 cycles v2 kits (Read 1: 28 cycles; Index Read (i7): 8 cycles; Read 2: 350 cycles) with 10% PhiX sequencing control v3 (Illumina).

#### QUANTIFICATION AND ANALYSIS

##### Preprocessing

For bulk experiments, raw Illumina paired-end reads of the CARLIN amplicon were merged using PEAR v0.9.11 (Zhang 2013) with parameters --min-overlap=1, --min-assembly-length=1, --min-trim-length=1, and --quality-threshold=30. Amplicon and transcriptome libraries prepared with 10X were preprocessed using CellRanger v2.2.0 (for 10X V2 libraries) and v3.0.2 (for 10X V3 libraries) according to the standardized workflow with default parameters. Transcriptome libraries were aligned against the mm10 reference genome provided by CellRanger.

Although no data presented in the paper was generated with InDrops (Zilionis 2017), it is a supported platform, though a modified version of the InDrops pipeline should be used (available at https://gitlab.com/hormozlab/indrops). The InDrops pipeline should be run with the parameters (LEADING=10, SLIDINGWINDOW=4:5, MINLEN=16) for Trimmomatic and (m=200, n=1, l=15, e=500) for Bowtie. Additionally, amplicon libraries prepared with InDrops should be processed with the '--no_clean_barcodes' flag, which preserves the uncleaned version of the Cell Barcode (CB) and logs QC information for the CB and UMI in the header (see *Filtering Reads* and *CB and UMI Error Correction* below).

Cells were additionally filtered by removing CBs in which the number of detected UMIs was below a threshold that was determined programmatically using MATLAB's findpeaks function based on the distribution of UMI counts (thresholds are listed in Supplementary Table 3). CBs where the percentage of UMIs corresponding to mitochondrial genes exceeded 15% were also discarded. The remaining CBs constitute a reference list against which CBs found in CARLIN amplicon sequencing can be collapsed (see *CB and UMI Error Correction*).

##### CARLIN Pipeline

Custom software was developed in MATLAB to perform alignment and allele calling, which can handle both bulk and single-cell amplicon sequencing of the CARLIN array. The software is trivial to install, simple to run, and produces data diagnostics useful for an experimentalist. The CARLIN software package is available at https://gitlab.com/hormozlab/carlin.

*Filtering Reads*

For bulk sequencing runs of genomic DNA, reads were filtered on possessing an exact match to the 20 bp CARLIN_fwd2 primer starting at the 11$^{th}$ bp of the read, since the UMI-tagging reaction adds a 10 bp randomer upstream of the CARLIN_fwd2 primer (Supplementary Table 1; Methods). To ensure that the entire CARLIN sequence was read, we retained reads that had good alignment to the 31 bp pre-polyA UTR sequence (hereafter referred to as the secondary sequence; Supplementary Table 1) located downstream of the 10th CARLIN target site – specifically, we require MATLAB's nwalign function with glocal=true (hereafter referred to as just nwalign) to return a score >=30.

For bulk sequencing runs of RNA, reads were reverse-complemented as the read is anti-sense. Because the UMI tagging protocol results in a 10 bp randomer being appended to the 20 bp CARLIN_rev primer (Supplementary Table 1; Methods), we retained reads that had an exact match to the primer, starting at 30 bp upstream of the end of the read. Additionally, to ensure that the whole CARLIN sequence was read and to simplify subsequent alignment, we only retained reads that had good alignment to the CARLIN_fwd2 primer and secondary sequence (nwalign scores >=15 and 30 respectively).

For all bulk reads, there is additional filtering to only retain reads with UMIs that have a QC of at least 30 at all 10 bps.

For sequencing runs of 10X CARLIN amplicon libraries, to ensure the whole CARLIN construct is sequenced, reads were filtered on possessing an exact match to the CARLIN_fwd2 primer starting at the 1$^{st}$ base pair, and a good alignment (nwalign scores >= 30) to the secondary sequence. Additionally, reads in which the CB or UMI are of the incorrect length, have uncalled base pairs, or in which the QC of any base pair is < 20, were discarded.

For all reads which pass the above filters, the CARLIN sequence was extracted by trimming the flanking primers and secondary sequence. If the resulting sequence has any uncalled base pair or was shorter than 26 bps (the combined length of the prefix, first conserved site and postfix – see Figure 1B), it was discarded.

*Alignment*

Here, we will describe the procedure used to determine how a CARLIN read has been altered with respect to the unmodified CARLIN sequence (referred to hereafter as the reference; see CARLIN reference sequence in Supplementary Table 1). As Cas9 modifications are expected to predominantly be indels, to identify alterations, we first aligned the CARLIN sequences against the reference. We found that existing alignment algorithms do not account for where Cas9 alterations are expected to appear along the reference (3 bp upstream of the PAM sequence, in a region referred to here as the cutsite). For example, NeedleAll (Rice 2000), a software package that implements the standard Needleman-Wunsch (NW) algorithm, was used in GESTALT (McKenna 2016), but yielded mixed results when aligning 75 modified CARLIN sequences read out using bulk Sanger sequencing. First, the NW algorithm did not preferentially select indel locations to coincide with the expected activity of Cas9 when multiple indel locations were equally plausible. Second, the default parameters used in the NW algorithm (NUC44 scoring matrix, gap opening penalty of 10, gap extension penalty of 0.5), precludes insertions and deletions from occurring consecutively, which is unrealistic given that the activity of Cas9 can result in such alterations. Third, the default value of the gap extension penalty parameter penalizes insertions and deletions according to length. Finally, customizing these parameters cannot cluster the alterations into expected regions of Cas9 activity.

To overcome these limitations, we developed our own alignment algorithm. Our algorithm is a modified version of the NW algorithm that performs alignment while accounting for where along the sequence alterations are expected with respect to the reference. We introduced a site-dependent cost function that penalizes alterations outside the expected regions of Cas9 activity. The cost function was minimized using a dynamic programming approach. Although the parameters used are specific to CARLIN, they can be modified to accommodate other systems that rely on Cas9 editing. Next, we describe our algorithm in full detail.

Let $\mathcal{N} = \{A, C, G, T\}$ be the set of nucleotides, $B$ denote a gap, and $\mathcal{N}_+ = \mathcal{N} \cup B$. Let $\vec{s} = [s_1 \dots s_j \dots s_J] \in \mathcal{N}^{1\times J}$, $J \geq 1$, be the nucleotide string of length $J$ to align and denote the reference of length $K$ by $\vec{r} = [r_1 \dots r_k \dots r_K] \in \mathcal{N}^{1\times K}$, $K \geq 1$. Let $\vec{s_j} \in \mathcal{N}^{1\times j}$, $0 \leq j \leq J$, and $\vec{r_k} \in \mathcal{N}^{1\times k}$, $0 \leq k \leq K$, be prefix strings of $\vec{s}$ and $\vec{r}$ ending with $s_j$ and $r_k$ respectively (or empty strings if $j = 0$ or $k = 0$). Denote by $Q\colon \mathcal{N}_+^{1\times L} \to \mathcal{N}^{1\times L'}, L \geq L'$, the operator which removes gaps from the input string.

Define $\vec{a}_{j,k} = \begin{bmatrix} \vec{a}_1 \\ \vec{a}_2 \end{bmatrix} \in {\mathcal{N}_+}^{2\times L}$, with $\max(j,k) \leq L \leq j+k$, as an alignment of $\vec{s_j}$ against $\vec{r_k}$ with $\vec{s_j} = Q(\vec{a}_1)$ and $\vec{r_k} = Q(\vec{a}_2)$, and let $V\colon {\mathcal{N}_+}^{2\times L} \to \mathbb{R}$ be a scoring function, that yields higher values for better alignments. We do not enumerate all alignments and rank them according to any explicit evaluation of this scoring function. Instead, we use a recursive formulation where we determine the maximum possible score for $\vec{a}_{j,k}$. This requires us to distinguish between three kinds of alignment: neither of the terminal characters are gaps, or the two cases where one terminal character is a gap but not the other. Using our notation, the three cases are (i) $\vec{m}_{j,k} = \begin{bmatrix} \vec{m}_1 \\ \vec{m}_2 \end{bmatrix} \in {\mathcal{N}_+}^{2\times L}$ in which $m_{1,L} \in \mathcal{N}$ and $m_{2,L} \in \mathcal{N}$, (ii) $\vec{d}_{j,k} = \begin{bmatrix} \vec{d}_1 \\ \vec{d}_2 \end{bmatrix} \in {\mathcal{N}_+}^{2\times L}$ in which $d_{1,L} = B$ and $d_{2,L} \in \mathcal{N}$, and (iii) $\vec{\imath}_{j,k} = \begin{bmatrix} \vec{\imath}_1 \\ \vec{\imath}_2 \end{bmatrix} \in {\mathcal{N}_+}^{2\times L}$ in which $i_{1,L} \in \mathcal{N}$ and $i_{2,L} = B$. Denote by $\mathcal{M}_{j,k} = \{\vec{m}_{j,k}\}$, $\mathcal{D}_{j,k} = \{\vec{d}_{j,k}\}$, $\mathcal{I}_{j,k} = \{\vec{\imath}_{j,k}\}$ the set of all such alignments.

We can then note the following recursive relationships, which simply state that an alignment of two strings can be decomposed into an alignment of two substrings, comprised of the first characters through to the penultimate characters, and an alignment of the terminal characters. We write down this recursive relationship for the three kinds of alignments defined above, using $\uplus$ to denote string concatenation.

$$
\begin{aligned}
\mathcal{M}_{j,k} = &\left\{\vec{m}_{j-1,k-1} \uplus \begin{bmatrix} s_j \\ r_k \end{bmatrix}, \vec{m}_{j-1,k-1} \in \mathcal{M}_{j-1,k-1}\right\} \\
\cup \quad &\left\{\vec{d}_{j-1,k-1} \uplus \begin{bmatrix} s_j \\ r_k \end{bmatrix}, \vec{d}_{j-1,k-1} \in \mathcal{D}_{j-1,k-1}\right\} \\
\cup \quad &\left\{\vec{\imath}_{j-1,k-1} \uplus \begin{bmatrix} s_j \\ r_k \end{bmatrix}, \vec{\imath}_{j-1,k-1} \in \mathcal{I}_{j-1,k-1}\right\} \\
\mathcal{D}_{j,k} = &\left\{\vec{m}_{j,k-1} \uplus \begin{bmatrix} B \\ r_k \end{bmatrix}, \vec{m}_{j,k-1} \in \mathcal{M}_{j,k-1}\right\} \\
\cup \quad &\left\{\vec{d}_{j,k-1} \uplus \begin{bmatrix} B \\ r_k \end{bmatrix}, \vec{d}_{j,k-1} \in \mathcal{D}_{j,k-1}\right\} \\
\cup \quad &\left\{\vec{\imath}_{j,k-1} \uplus \begin{bmatrix} B \\ r_k \end{bmatrix}, \vec{\imath}_{j,k-1} \in \mathcal{I}_{j,k-1}\right\}
\end{aligned}
$$

$$\mathcal{I}_{j,k} = \left\{\vec{m}_{j-1,k} \uplus \begin{bmatrix} s_j \\ B \end{bmatrix}, m_{j-1,k} \in \mathcal{M}_{j-1,k}\right\}$$
$$\cup \quad \left\{\vec{d}_{j-1,k} \uplus \begin{bmatrix} s_j \\ B \end{bmatrix}, \vec{d}_{j-1,k} \in \mathcal{D}_{j-1,k}\right\}$$
$$\cup \quad \left\{\vec{\imath}_{j-1,k} \uplus \begin{bmatrix} s_j \\ B \end{bmatrix}, \vec{\imath}_{j-1,k} \in \mathcal{I}_{j-1,k}\right\}$$

The base cases are the empty sets $\mathcal{M}_{j\geq 0,0} = \mathcal{M}_{0,k\geq 0} = \mathcal{D}_{j\geq 0,0} = \mathcal{I}_{0,k\geq 0} = \emptyset$, and the singleton sets $\mathcal{D}_{0,k} = \vec{d}_{0,k}$ for $1 \leq k \leq K$ and $\mathcal{I}_{j,0} = \vec{\imath}_{j,0}$ for $1 \leq j \leq J$.

We now elucidate the score to rank the alignments in these sets. Briefly, we penalize deletions that do not begin and finish near the expected cutsites, and insertions that do not occur near the expected cutsites. We do not favor insertions over deletions, discriminate based on the length of the alteration, or penalize consecutive deletions or insertions.

Let $C_M: \mathcal{N} \times \mathcal{N} \rightarrow \mathbb{R}$, be a scoring function for aligning different nucleotides (we use the same NUC44 scoring matrix used in the default NW algorithm). Define $P_{D,B}(k) \geq 0$ and $P_{D,E}(k) \geq 0$, $1 \leq k \leq K$ to be site-dependent penalties for beginning and ending a deletion at $r_k$ from alignments which end or begin in paired nucleotides respectively. Similarly, define $P_{I,B}(k) \geq 0$, and $P_{I,E}(k) \geq 0$ for $1 \leq k \leq K$ to be the site-dependent penalty for beginning an insertion after $r_k$ and ending an insertion before $r_k$ respectively. Let $M_{j,k}$, $D_{j,k}$, $I_{j,k}$ be the highest scores of the alignments in $\mathcal{M}_{j,k}$, $\mathcal{D}_{j,k}$ and $\mathcal{I}_{j,k}$ respectively.

$$M_{j,k} = \max_{\vec{m}_{j,k} \in \mathcal{M}_{j,k}} V(\vec{m}_{j,k})$$
$$D_{j,k} = \max_{\vec{d}_{j,k} \in \mathcal{D}_{j,k}} V(\vec{d}_{j,k})$$
$$I_{j,k} = \max_{\vec{\imath}_{j,k} \in \mathcal{I}_{j,k}} V(\vec{\imath}_{j,k})$$

Using the set recurrences above and the introduced scoring scheme, we arrive at a new set of scalar recurrences for the highest scores, so that we don't have to compute these quantities explicitly by exhaustively searching over a full enumeration of alignments.

$$M_{j,k} = \max\{M_{j-1,k-1}, D_{j-1,k-1} - P_{D,E}(k-1), I_{j-1,k-1} - P_{I,E}(k)\} + C_M(s_j, r_k)$$
$$D_{j,k} = \max\{M_{j,k-1} - P_{D,B}(k), D_{j,k-1}, I_{j,k-1}\}$$
$$I_{j,k} = \max\{M_{j-1,k} - P_{I,B}(k), D_{j-1,k}, I_{j-1,k}\}$$

As evident in the last two equations, there is no incremental cost in extending an insertion or deletion along either one of the two strings. Since insertions and deletions are penalized equally, we make the following simplifying assumption, which in turn also reduces the number of parameters: $P_{D,B}(k) = P_{I,B}(k) \equiv P_B(k)$ and $P_{D,E}(k) = P_{I,E}(k) \equiv P_E(k)$. We abuse notation to let $P_B(0)$ be the penalty for starting insertions prior to the first character of the reference. Consistent with the previously defined base cases, the initialization is $M_{j>0,0} = M_{0,k>0} = D_{j\geq 0,0} = I_{0,k\geq 0} = -\infty$, $D_{0,k>0} = P_B(1)$, $I_{j>0,0} = P_B(0)$, $M_{0,0} = 0$.

There is no guarantee that a single alignment $\vec{a}_{j,k}$ achieves the highest score $(\max\{M_{j,k}, D_{j,k}, I_{j,k}\})$. Multiple alignments can achieve the maximum score within each of $\mathcal{M}_{j,k}$, $\mathcal{D}_{j,k}$ and $\mathcal{I}_{j,k}$, and also across the three. To select the optimal alignment amongst the alignments that share the maximum score, we need to retain which argument in the recurrence relationships for $\mathcal{M}_{j,k}$, $\mathcal{D}_{j,k}$ and $\mathcal{I}_{j,k}$ (see above) realize the maximum score. Let $E : \mathcal{N}_+^{2\times L} \to \mathcal{N}_+^{2\times(L-1)}$, $L > 0$, be the operator which strips the terminal character from the aligned sequence and reference. Given $S, \mathcal{S} \in \{\mathcal{M}, \mathcal{D}, \mathcal{I}\}$, let $R_{j,k}^{(S,\mathcal{S})}$, $0 \le j \le J$, $0 \le k \le K$, be an indicator function such that

$$R_{j,k}^{(S,\mathcal{S})} = \begin{cases} 1, & \text{if } \exists\ \vec{a}_{j,k} \in \underset{\vec{b}_{j,k} \in \mathcal{S}_{j,k}}{\operatorname{argmax}} V(\vec{b}_{j,k}) \text{ such that } E(\vec{a}_{j,k}) \in S \\ 0, & \text{otherwise} \end{cases}$$

For example, if both the first and third terms realize the maximum in the recurrence for $D_{j,k}$, then we set $R_{j,k}^{(\mathcal{M},\mathcal{D})} = R_{j,k}^{(\mathcal{I},\mathcal{D})} = 1$ and $R_{j,k}^{(\mathcal{D},\mathcal{D})} = 0$. We will use this indicator function to select the optimal alignment as described below. In accordance with the previous base cases, the initialization is $R_{j\ge0,0}^{(\cdot,\mathcal{M})} = R_{0,k\ge0}^{(\cdot,\mathcal{M})} = R_{j\ge0,0}^{(\cdot,\mathcal{D})} = R_{0,k\ge0}^{(\cdot,\mathcal{I})} = 0,\ R_{0,k>0}^{(\mathcal{D},\mathcal{D})} = R_{j>0,0}^{(\mathcal{I},\mathcal{I})} = 1$.

The alignment algorithm is implemented in two stages, a forward stage in which we tabulate the maximum score, and a reverse stage in which we construct the optimal alignment. In the forward stage, the maximum scores are first initialized as outlined above. Next, we solve the recurrences using a dynamic programming approach in $O(JK)$ time by sweeping across all nucleotides in the sequence from $j = 1$ to $J$, and for each value of $j$, iterating over all the nucleotides in the reference, $k = 1$ to $K$. At each $(j, k)$ ordered pair, we compute and store $M_{j,k}$, $D_{j,k}$ and $I_{j,k}$ in three two-dimensional arrays of size $(J + 1) \times (K + 1)$, and update a $3 \times 3 \times (J + 1) \times (K + 1)$ Boolean array for $R_{j,k}^{(S,\mathcal{S})}$, which retains the alignment type (gap in the reference, gap in the sequence, or no gap) that achieves the maximum scores.

In the reverse stage, we incrementally build the optimal alignment $\vec{a}$ by backtracking from the last aligned pair of nucleotides $(J, K)$ using the following rules, picking the first lettered rule that applies within each numbered rule. These rules were formulated to ensure continuity among mutation types, which allows us to group distinct mutations more easily (see *Mutation Calling*) and attribute them to a single Cas9 cutting event.

1. Let $j = J$, $k = K$, $\vec{a} \in \mathcal{N}_+{}^{2\times0}$ .
    a. If $M_{J,K} \ge \max\{D_{J,K}, I_{J,K}\}$, let $\mathcal{S} = \mathcal{M}$.
    b. If $D_{J,K} \ge \max\{M_{J,K}, I_{J,K}\}$, let $\mathcal{S} = \mathcal{D}$.
    c. If $I_{J,K} \ge \max\{M_{J,K}, D_{J,K}\}$, let $\mathcal{S} = \mathcal{I}$.

2. Repeat rules 3-5 until $j = 0$ and $k = 0$.

3. If $\mathcal{S} = \mathcal{M}$:

a. If $R_{j,k}^{(\mathcal{M},\mathcal{M})} = 1$, let $\mathcal{S} = \mathcal{M}$.
b. If $R_{j,k}^{(\mathcal{D},\mathcal{M})} = 1$, let $\mathcal{S} = \mathcal{D}$.
c. If $R_{j,k}^{(\mathcal{I},\mathcal{M})} = 1$, let $\mathcal{S} = \mathcal{I}$.

Let $\vec{a} \leftarrow \begin{bmatrix} s_j \\ r_k \end{bmatrix} \uplus \vec{a}$, $j \leftarrow j-1$, $k \leftarrow k-1$.

4. If $\mathcal{S} = \mathcal{D}$:
   a. If $R_{j,k}^{(\mathcal{D},\mathcal{D})} = 1$, let $\mathcal{S} = \mathcal{D}$.
   b. If $R_{j,k}^{(\mathcal{M},\mathcal{D})} = 1$, let $\mathcal{S} = \mathcal{M}$.
   c. If $R_{j,k}^{(\mathcal{I},\mathcal{D})} = 1$, let $\mathcal{S} = \mathcal{I}$.

   Let $\vec{a} \leftarrow \begin{bmatrix} B \\ r_k \end{bmatrix} \uplus \vec{a}$, $k \leftarrow k-1$.

5. If $\mathcal{S} = \mathcal{I}$:
   a. If $R_{j,k}^{(\mathcal{I},\mathcal{I})} = 1$, let $\mathcal{S} = \mathcal{I}$.
   b. If $R_{j,k}^{(\mathcal{M},\mathcal{I})} = 1$, let $\mathcal{S} = \mathcal{M}$.
   c. If $R_{j,k}^{(\mathcal{D},\mathcal{I})} = 1$, let $\mathcal{S} = \mathcal{D}$.

   Let $\vec{a} \leftarrow \begin{bmatrix} s_j \\ B \end{bmatrix} \uplus \vec{a}$, $j \leftarrow j-1$.

The result is a unique alignment $\vec{a}$ such that $V(\vec{a}) = \max\{M_{J,K}, D_{J,K}, I_{J,K}\}$.The algorithm above can be used to generate improved alignments for any CRISPR-Cas9 system for suitably chosen penalties $P_B(k)$ and $P_E(k)$.

Lastly, we turn our attention to the values chosen for the penalty functions, for CARLIN specifically. Because we expect the alterations to be localized to the cutsites, the minimum value of the penalty was set to occur within 3 bp upstream of the PAM sequences with the penalty increasing linearly moving away from this location. We used the same penalty function for all the target sites. We manually selected the parameters of the penalty function, i.e. the minimum value and the slopes, to promote alignments comprised of insertions and deletions in the cutsites for the 75 CARLIN sequences characterized using bulk Sanger sequencing. The same parameters also resulted in desired alignment properties when applied to the bulk RNA sequencing data used to make the allele bank (Supplementary Figure 2A-C). For the numerical values of the penalty functions, refer to Supplementary Table 2.

*Motif Classification*

We partitioned the reference into 31 consecutive regions (called motifs), according to the locations of the reference prefix sequence, conserved sites, cutsites, PAM+linkers, and postfix sequence (Figure 1B). After aligning a sequence to the reference, we identified the same motifs in the aligned sequence according to the nucleotide boundaries of the motif in the reference. Each aligned motif was then assigned a classification: (1) ‘N’: all the nucleotides of the sequence match that of the reference exactly and there are no gaps in the sequence or reference. (2) ‘E’: motif is completely absent in the aligned sequence. (3) ‘D’: any (but not all) bps of the motif are deleted in the sequence, and there are no gaps in the reference. (4) ‘M’: there are no insertions or deletions relative to the reference, only substitutions. (5) ‘I’ otherwise. These motifs were used to correct potential sequencing errors and establish a consensus sequence across multiple reads (as described below).

*Sequence Error Correction*

Sequencing and amplification (technical) errors can, in addition to Cas9 activity, also introduce alterations in the CARLIN sequence. We assumed that any individual SNP that occurs outside of a 3 bp window around the cutsite is a technical error. To remove these technical errors, we took the aligned CARLIN sequences and reverted the SNPs in these regions (classified as ‘M’ in non-cutsite motifs) to the corresponding base pair in the reference. We also trimmed nucleotides in the aligned sequence that extended beyond the reference in either direction.

*CB and UMI Error Correction*

CBs and UMIs are also subject to the same technical errors as the CARLIN sequence. Denoising of CBs and UMIs is performed according to the directional adjacency method developed in UMI-tools (Smith 2017). In short, the method starts by making a list of tags (either CBs or UMIs) sorted in descending order by the number of reads corresponding to each tag. Next, a directed graph is constructed with nodes representing tags. The objective is to connect the nodes whose tags are different only due to technical errors.

To do so, first, the top-ranked node is selected as the current node. An edge is drawn from the current node to all candidate nodes that satisfy the following criteria: the candidate node does not already have an incident edge, the tag of the candidate node is different by only one base pair from the tag of the current node, and the number of reads of the candidate node’s tag is less than half of the current node’s tag. The current node is then updated to be the next most common node. The process is repeated until the entire sorted list has been traversed, resulting in disjoint sets of connected nodes (connected components). In each connected component, the reads of all the tags are merged under the tag of the top-ranked node in that connected component.

For bulk samples where no CB exists, we do not apply this algorithm directly to all of the detected UMIs. Instead, we first group the UMIs by their consensus CARLIN sequence with no read threshold (see *Consensus Calling* and *Read Thresholds for Denoised CBs and UMIs* below for definition of consensus sequence and use of read thresholds respectively). Next, the directional adjacency method (describe in the previous paragraph) is applied across all UMIs with the same consensus sequence. This extra step prevents accidental merging of UMIs that differ by only 1 bp when the number of detected UMIs is large with respect to the possible UMI diversity.

For single-cell data, directional adjacency denoising is also used but with CBs for tags. If there is a reference CB list (for e.g. obtained after preprocessing of the transcriptome library, see *Preprocessing*), then we include two additional requirements: (i) top-ranked nodes in each connected component need to belong to the reference CB list (or else the whole connected component is discarded), and (ii) candidate nodes cannot belong to the reference CB list. Next, for each denoised CB (which is now guaranteed to be in the reference list, if supplied), directional adjacency denoising is performed on its constituent UMIs. The result of this procedure is a denoised list of CBs and/or UMIs.

*Read Thresholds for Denoised CBs and UMIs*

Since we do not necessarily have a ground truth list of tags which we expect to detect in the data, tags with few reads may be spurious, even if they have been denoised. Additionally, to call a consensus sequence for each tag, we need to be able to find a common sequence among a sufficiently large number of reads; repeat occurrences of a CARLIN sequence over many reads gives us confidence that the sequence is the correct one. Averaging over many reads also allows us a secondary mechanism of correcting for technical errors (especially at cutsites, which we left untouched in the *Sequence Error Correction* section). Here we describe how we determine a threshold on what is a sufficient number of reads.

Let $R_i$ be a sorted list of the number of reads associated with each of $N$ denoised tags (UMIs/CBs) with $R_1 \geq \ \dots \geq R_N$. We only attempt to call a consensus sequence for tag $i$ if $R_i$ meets a minimum threshold, $T$, for the number of reads, and discard tags which fail to achieve this threshold.

$$T = \max\{\left\lceil\frac{R_{[0.01N]}}{10}\right\rceil, R_{N_{expected}}, \left\lceil\frac{\sum_i R_i}{N_{expected}}\right\rceil, \lceil R_1 p(1-p)^{L-1}\rceil, 10\}$$

$N_{expected}$ is the expected number of tags in the experiment (or $\infty$ if unknown or $N_{expected} > N$, so that $R_\infty = 0$), $p$ is the 95$^{th}$ percentile value of the transformed QC scores (error probability) across all tag bps for all filtered reads, and $L$ is the number of base pairs in the tag.

Our threshold function is a heuristic and was empirically optimized by comparing the length distribution of the consensus alignments called from the consensus sequence (see *Consensus Calling*) to fragment length analysis of the same libraries (Supplementary Figure 2D). We observed that the log-log plot of the rank-ordered number of reads across the tags, $R_i$, exhibited a plateau (roughly constant number of reads) after the first percentile with a sharp fall off (a knee) when the number of reads decreased to one tenth of that of the plateau. Similar empirical criteria are used to select single-cell barcodes by other single-cell analysis tools such as CellRanger. The next two terms of the threshold function ensure that the number of tags does not exceed the expected number of molecules. The fourth term sets a conservative estimate on the number of reads expected from a single sequencing error across a tag of length L. Finally, we also required that any tag is observed in at least 10 reads. This last condition reflects the minimum number of sequences we would like to have for consensus calling (see below). The choices for the threshold function were made conservatively to minimize false-positives.

For SC data for which a reference CB list is not available, first $R_i$ is tabulated as the number of reads for a given CB. A CB read threshold, $T_{CB}$, is computed using $R_i$ as described above. Next, the number of reads is also tabulated for each CB-UMI pair, to compute $T_{UMI}$. Only cell barcodes which have at least $T_{CB}$ reads, and within each CB, UMIs which have at least $T_{UMI}$ reads are retained.

For SC data for which a reference CB list is available, only CBs found in the reference CB list survive the denoising procedure. Since the reference CB list is separately subject to quality control in the software generating the list (for e.g. CellRanger), we trust all denoised CBs, so that the threshold is only required to ensure there are sufficient reads for consensus calling. To that end, we only use the last term in our heuristic function, effectively setting $T_{CB} = T_{UMI} = 10$. The mean number of reads, $R_i$, among CBs that pass this threshold criterion is typically one or two orders of magnitude larger than $T$ (see Supplementary Table 3).

*Consensus Calling*

Next, we describe the procedure used to call the consensus sequence for tags that pass the threshold criterion. First, we selected the tags for which greater than 50% of the reads are of the same length and possess the same 31-character motif classification string (see *Motif Classification* above). Next, for the selected tags, we retained the reads that comprised this majority. The consensus sequence was constructed bp-by-bp by selecting the most commonly occurring nucleotide across all the retained reads (i.e. taking a per bp mode of the retained reads).

For single-cell data, this procedure is performed on each UMI of a CB separately. Next, we collected the consensus sequence across different UMIs that passed the threshold and selection criteria. If more than 50% of the consensus sequences are of the same length and possess the same 31-character motif classification string, we proceeded as follows. We retained the consensus sequences that comprised this majority. The CB consensus sequence was constructed bp-by-bp, selecting the most commonly occurring nucleotide across the retained UMI consensus sequences.

To construct the consensus alignment, we removed the gaps from the consensus sequence and realigned to the reference. To define CARLIN alleles, we pooled all consensus alignments, and removed all repeated instances, so that each consensus alignment is represented only once. We define an allele to be an element of the resulting set, so that alleles are unique by definition. An allele is defined to be edited if its consensus alignment does not match the reference exactly. In the text, edited transcript/cell refers to a UMI/CB which has a corresponding CARLIN allele that is edited. The frequency of an allele in a sample is the number of tags whose consensus alignment matches that allele.

*Mutation Calling*

To explicitly associate mutations with the alterations harbored by an allele, we first tabulated the starting and ending locations of the deletions, insertions, and substitutions in the consensus alignment. Next, to account for a single Cas9 cutting event that could result in multiple alterations, we combined two mutations if they were adjacent – the ending location of one mutation was one bp upstream of the starting location of the other— or if the ending location of one was co-localized to the same cutsite as the starting location of the other. We iterated the process of combining mutations until no further combinations were possible. The combined mutations were always designated as indels. For example, a typical allele can harbor a single bp deletion, and a 5 bp insertion at the cutsite of target site 3, two mutations which are merged into a single indel event using this procedure.

For the purpose of preparing figures, we considered all substitutions as indels (1 bp deletion of the reference nucleotide, and 1 bp insertion of the mismatched sequence nucleotide). In figures where only insertions and deletions are shown (e.g. Figure 1C,D,F), indels are represented twice, once as an insertion corresponding to the length of the new sequence, and once as a deletion for the length of the deleted portion of the reference.

*CARLIN Potential*

To quantify the extent to which a CARLIN allele is altered with respect to the reference, we define the CARLIN potential of an allele as the number of target sites whose sequence and the sequence of the adjacent PAM+linker domain exactly matches that of the reference. Alterations in the prefix and postfix are considered to be alterations of the first and last target site respectively. The CARLIN potential is a whole number between 0 and 10.

**Quantifying Diversity**

*Effective Alleles*

Since independent Cas9 activity across different cells may lead to the same altered sequences, the number of alleles detected will generally be less than the number of cells, even if all cells are edited. Furthermore, not all alleles are equally likely. Some occur more frequently than others.

Ideally, we want detection of a particular allele to restrict the cell in which the allele is observed to the smallest possible subset of the original population. For a given number of alleles, this occurs if all alleles are equally likely (i.e. the observation carries maximal information in the language of information theory). For the case where the allele frequencies are not equal, the observation of a particular allele is, on average, not as informative for constraining the subpopulation of cells from which the allele came. How many equally likely alleles are required to carry the same amount of information in this case?

To answer this question, we devised a metric, the effective number of alleles, calculated as $2^H$, where $H$ is the Shannon entropy of the normalized allele frequency distribution across the edited cells. In the limit that each edited allele appears in an equal number of cells, the effective number of alleles is simply the number of edited alleles. Conversely, in the limit where a single allele appears in all edited cells, the number of effective alleles is 1. The effective allele measure seeks to discount the effect of over-represented alleles, which are less informative. The effective number of alleles in the bank of 44000 alleles is 14700.

*Diversity Index*

The diversity index is computed by dividing the effective number of alleles (see above) by the number of cells in the population. (Alternatively, dividing by the number of edited cells can be used to compute diversity index across only the edited cells, independent of the fraction of cells edited). The diversity index is 0 when only the reference is present in the sample, and 1 when the reference is entirely absent and each cell has its own allele. The diversity index is $\alpha$ when the cutting efficiency is $\alpha$, and each edited cell has a distinct allele, and is less than $\alpha$ if some cells share the same allele. The reciprocal of the diversity index is the average number of cells labeled by each effective allele.

*Allele Bank*

To estimate the total number of alleles which can be generated by the CARLIN system, we pooled the edited transcripts collected from granulocytes harvested from 3 mice, to create a bank of ~233K transcripts resulting in ~32K alleles. As granulocytes are not expected to proliferate appreciably in the 3 days between induction and collection, shared alleles across multiple cells are coincidental and not due to shared ancestry of the cells.

*Estimating Number of Unseen Alleles*

The total number of alleles that CARLIN can generate is the sum of the number of alleles observed in the bank and the number of alleles that have gone undetected because only a finite number of cells were measured. To extrapolate the total number of alleles, we need to estimate the number of unobserved alleles. To intuitively understand the extrapolation, we consider two extreme cases. If most alleles are only observed once, then we expect many new alleles would turn up under more exhaustive sampling. (In the limit that all alleles are observed only once, the total number of alleles is indeterminate). Conversely, if most alleles are observed many times, the system has been sufficiently sampled and few unobserved alleles are expected. To estimate the number of unobserved alleles, we use tools developed in the field of ecology, where the problem is referred to as the unseen species problem. See (J. A. Bunge 2014) for example, for a review of techniques in the field.

*Non-Parametric Diversity Estimation*

We use the Smoothed Goodman-Toulmin estimator (SGT) proposed by (Orlitsky 2016) to estimate the expected number of alleles, $N$, in $M$ observations given that $n$ alleles were detected in $m$ observations $(m < M)$. This estimator is near-optimal in the sense of mean-squared error and works for $M$ of the order of $O(m\log m)$, the theoretical extrapolation limit (Orlitsky 2016). To check the consistency of the estimator, we subsample the data to have $\tilde{m} \leq m$ observations (colored dots in Figure 3G), and compute the expected number of alleles for an extrapolated number of observations in the interval $\widetilde{m} \leq \widetilde{M} \leq \widetilde{m}\log\widetilde{m}$ (dotted lines in Figure 3G colored to correspond to the subsample size).

*Parametric Diversity Estimation*

To validate the non-parametric method, and obtain an analytic expression for occurrence frequency of the alleles for subsequent calculations, we tabulate the frequency of allele $i$ as $X_i$ and generate a histogram of frequency counts, $h(Z) = \sum_i \delta_{Z,X_i}$, representing the number of alleles that are detected $Z$ times. Note that $\sum_Z h(Z) = n$, the number of edited alleles observed, and $\sum_Z Zh(Z) = \sum_i X_i = m$, the number of observations of edited alleles.

Conceptually, sampling of an allele can be approximated as a Poisson process. The number of observations of a particular allele, $X_i$, given a total number of observations, $M$, is drawn from a Poisson distribution with a rate parameter, $\lambda_i$, that is proportional to the prevalence of that allele in the bank. The Poisson approximation allows for an elegant analytical solution for estimating the occurrence frequency of an allele:

$$P(X_i = k; \lambda_i, M) = \frac{(\lambda_i M)^k}{k!} e^{-\lambda_i M}$$

To model the variability in the prevalence of different alleles, we need to associate a Poisson rate parameter with each allele. Although in general, each allele can have a unique Poisson rate parameter, for simplicity, we assume that the rate parameters are drawn from a distribution that can be succinctly described using a few parameters.

We use CatchAll (v4.0), a software package described in (J. L. Bunge 2012), to fit a distribution of intensive Poisson rate parameters for alleles, $f_\theta(\lambda)$, to our observed data. If all alleles are equally prevalent with $\lambda = \hat{\lambda}$, $f_\theta(\lambda) = \delta(\lambda - \hat{\lambda})$, and $h(Z)$, when normalized, would converge to a Poisson distribution. CatchAll determined that for our data, $h(Z)$ mostly closely resembles a mixture geometric distribution so that $f_\theta(\lambda)$ is a 4-component exponential mixture model.

The probability that an allele remains unobserved after $m$ observations is then given by

$$P_0 = P(X = 0; m) = \int P(X = 0|\, \lambda; m) P(\lambda) d\lambda = \int e^{-\lambda m} f_\theta(\lambda) d\lambda$$

The number of unseen alleles is therefore $nP_0/(1 - P_0)$ and the total number of alleles is given by $N = n/(1 - P_0)$. For our bank, $m \approx 233000$, $n \approx 32000$ and $P_0 \approx 0.28$ so that the number of unseen alleles is estimated to be $\approx 12000$ and consequently, the total number of alleles is estimated to be $N \approx 44000$. The curves in Figure 3H are obtained by using this estimate for $N$, and the endpoints of the 95% CI of $N$, in the estimators for interpolation and extrapolation provided in equations (6) and (12) respectively of (Colwell 2012).

*Prevalence of Unseen Alleles*

Given our analytical model, we can now determine the distribution of the Poisson rate parameters for the unobserved alleles. It is a conditional distribution given by Bayes' rule:

$$P(\lambda|X=0;m)=\frac{P(X=0|\lambda;m)P(\lambda)}{P(X=0;m)}$$

For simplicity, we assign all alleles unobserved in the bank the mean Poisson rate under this conditional distribution.

$$\lambda_0=\mathbb{E}_{P(\lambda|X=0;m)}[\lambda]$$

*Usage of Allele Rates*

For subsequent experiments, we query the alleles observed in the experiment against the alleles in the bank. If an observed allele matches an allele in the bank, that allele is assigned the rate parameter $\lambda_i=\frac{X_i}{m}$ based on its empirical prevalence in the bank, where $X_i$ is the frequency of the allele in the bank and $m$ is the total number of observations constituting the bank. Otherwise, we assign it the rate $\lambda_0$.

**Statistical Analysis**

*Limitations of a Finite Number of Alleles for Uniquely Marking Clones*

Here, we consider the implications of having a finite set of alleles with which we can mark a (possibly infinite) number of cells. How useful are the alleles in uniquely marking clones if multiple cells can be marked with the same allele? To answer this question, the following factors need to be considered: the frequency of the allele in the bank, the number of cells marked initially, potential proliferation of the cells after they are marked, and the number of cells sampled at the final timepoint.

Suppose that at induction, $C\geq 1$ cells are randomly assigned one of $N$ alleles, according to a distribution $\rho$ of allele probabilities. Let $L_j\in\{1\,\ldots.N\}$ be the allele label of the $j^{th}$ induced cell and let $C_i=\sum_{j=1}^{C}\delta_{L_j,i}$ be the number of cells marked with allele $i$. Then $P(C_i=c;C)=\binom{C}{c}\rho_i^c(1-\rho_i)^{C-c}$ and the conditional probability given that allele $i$ is present in at least one cell is

$$P(C_i=c|C_i>0;C)=\frac{P(C_i=c;C)}{1-P(C_i=0;C)}$$

We define the singleton probability and duplication probabilities as

$$p_{singleton}(i;C)=P(C_i=1|C_i>0;C)$$

$$p_{duplicate}(i;C) = P(C_i > 1 | C_i > 0; C) = 1 - p_{singleton}(i;C)$$

See Supplementary Figure 3B for an illustration of how $p_{singleton}$, the probability that an allele uniquely marks a cell, depends on $C$ and $N$ in the case that $\rho$ is uniform.

In practice, we do not know how the cells are labeled initially but instead observe alleles from an unbiased sampling of $M \geq 1$ cells at some later time. Subsequent to induction, assume that cell $j$ expands such that at the sampling time, the probability that cell $j$ is a progenitor of a sampled cell is given by $p_{progenitor}(j;b)$, where $b$ is a parameter, which will be defined later.

Our goal is to label each progenitor uniquely but it could be that some progenitors were coincidentally marked with the same allele. At the time of sampling, if we observe the same allele across multiple cells, we would like to know the probability that all the cells came from a single progenitor.

If we assume that the proliferation rates are not dependent on the allele label, we will be able to develop a useful bound on the above probability. Let $X_i$ be the number of cells in the observed sample marked with allele $i$. If progenitor proliferation rates are uncorrelated with progenitor allele labels, it follows that the distribution of allele frequencies in the sample is equal to that of the induced pool

$$\begin{aligned} P(X_i = x; M) &= \binom{M}{x}\left(\sum_{j=1}^{C} p_{progenitor}(j;b)\,\rho_i\right)^{x}\left(1 - \sum_{j=1}^{C} p_{progenitor}(j;b)\,\rho_i\right)^{M-x} \\ &= \binom{M}{x}\rho_i^x(1-\rho_i)^{M-x} \\ &= P(C_i = x; M) \end{aligned}$$

Let $Y_i$ be the number of progenitors of the sampled cells marked with allele $i$ (note the distinction from $C_i$, the number of progenitors marked with allele $i$). We know that generally $P(Y_i = m; M)$ depends on the induction itself (since $P(Y_i > C_i; M) = 0$, for example, when the realization of $C_i$ corresponds to the same experiment as $Y_i$). But what can we say about $P(Y_i = m; M)$ when we don't know $C_i$ for the same experiment, only $P(C_i = c; C)$?

First note that the conditional probability of the number of progenitors $Y_i$ given that allele $i$ is observed at least once is

$$\begin{aligned} P(Y_i = m | X_i > 0; M) &= \frac{P(Y_i = m; M)}{1 - P(X_i = 0; M)} \\ &= \frac{\sum_{k=0}^{M} P(Y_i = m | X_i = k) P(X_i = k; M)}{1 - P(X_i = 0; M)} \\ &= \frac{\sum_{k=m}^{M} P(Y_i = m | X_i = k) P(X_i = k; M)}{1 - P(X_i = 0; M)} \end{aligned}$$

The specific form of $P(Y_i = m|X_i = k)$ depends on the proliferation model. For an observed allele $i$, let us consider the ratio of the probability that more than one progenitor gave rise to the cells labeled with allele $i$ at the sampling timepoint, to the probability that more than one cell was labeled with allele $i$ at the initial timepoint:

$$\gamma\left(f = \frac{M}{C}\right) = \frac{P(Y_i > 1|X_i > 0; M)}{P(C_i > 1|C_i > 0; C)}$$

We note that we calculate these probabilities independently because in general we might not know the number of cells that were initially labeled with allele $i$. We show that $\gamma(1) \leq 1$, so that the probability of more than one progenitor giving rise to cells labeled with allele $i$ is smaller than the probability of labeling more than one cell with allele $i$ at the initial timepoint, given that the number of cells sampled is less than or equal to the initial number of cells:

$$\begin{aligned}
\gamma(1) &= \frac{P(Y_i > 1|X_i > 0; M)}{P(C_i > 1|C_i > 0; M)} \\
&= \frac{P(C_i > 0; M)}{P(X_i > 0; M)} \frac{P(Y_i > 1, X_i > 0; M)}{P(C_i > 1, C_i > 0; M)} \\
&= \frac{P(Y_i > 1; M)}{P(C_i > 1; M)} \\
&= \frac{\sum_{m'=2}^{M} P(Y_i = m'; M)}{\sum_{m'=2}^{M} P(C_i = m'; M)} \\
&= \frac{\sum_{m'=2}^{M} \sum_{k=m'}^{M} P(Y_i = m'|X_i = k) P(X_i = k; M)}{\sum_{m'=2}^{M} P(C_i = m'; M)} \\
&= \frac{\sum_{k=2}^{M} P(X_i = k; M) \sum_{m'=2}^{k} P(Y_i = m'|X_i = k)}{\sum_{m'=2}^{M} P(C_i = m'; M)} \\
&= \frac{\sum_{k=2}^{M} P(X_i = k; M) \left[1 - P(Y_i = 1|X_i = k)\right]}{\sum_{m'=2}^{M} P(C_i = m'; M)} \\
&\leq \frac{\sum_{k=2}^{M} P(X_i = k; M)}{\sum_{m'=2}^{M} P(C_i = m'; M)} \\
&= 1
\end{aligned}$$

This result can be generalized to bound any number of progenitor cells:

$$\frac{P(Y_i > m|X_i > 0; M)}{P(C_i > m|C_i > 0; C)} \leq 1, M \leq C$$

How close is $\gamma(1)$ to 1 for a given choice of parameters? Intuitively, the more heterogeneous the proliferation rates, the less likely we are to observe cells that share the same allele but not the same progenitor, since the clone that proliferates faster will tend to dominate the sample.

To show this mathematically we note $\gamma(1)$ is maximized when $P(Y_i = 1|X_i = k)$ is minimized for each $1 < k \leq M$. To determine when $P(Y_i = 1|X_i = k)$ is minimized, we turn our attention to the $b$ parameter of $p_{progenitor}(j;b)$. Let $b$ be any parameterization of $p_{progenitor}$ such that $p_{progenitor}(j;0) = \frac{1}{C}$, $\|p_{progenitor}(b)\|_\infty \to 1$ as $b \to \infty$, and $\frac{d\|p_{progenitor}(b)\|_\infty}{db} > 0$. Note that

$$P(Y_i = 1|X_i = k) = \sum_{j,L_j=i} \left( \frac{p_{progenitor}(j;b)}{\sum_{j,L_j=i} p_{progenitor}(j;b)} \right)^k$$

This is minimized for ∀ $k$ (for example, by Lagrange multipliers) when each of its constituent terms is $\frac{1}{C_i}$, so that $p_{progenitor}(j;b) = \frac{1}{C}$ i.e. $b = 0$.

To consider the effect of sampling, we resort to numerical simulations and consider a specific proliferation model given by $p_{progenitor}(j;b) = \frac{1}{Z(b)}\left(1 - \frac{j-1}{C-1}\right)^b$, which results in $p_{progenitor}(j;b) \geq p_{progenitor}(j+1;b)$ without loss of generality. $Z(b) = \sum_j p_{progenitor}(j;b)$ is a normalizing constant. We simulate induction of an initial pool with $N = 2000$ alleles (the results are insensitive to $N$) drawn from a uniform $\rho$, subsequent expansion using this proliferation model, followed by sampling of a different number of cells at the final timepoint (resulting in different instances of $f$). Supplementary Figure 3B shows $\gamma(f) \leq 1$ for different values of $f$ and $b$ where each data point is the mean of a 1000 simulations. The more uneven the proliferation rates (larger values of $b$), the less likely it is to observe the same allele shared across multiple cells that arise from different progenitors. However, as the number of observed cells increases (larger values of $f$), even progeny from slowly-proliferating progenitors that were redundantly marked are more likely to be observed.

In summary, if we assume that proliferation rates are not dependent on allele marking then:

1. The probability that cells that share a given allele at the final timepoint came from more than one progenitor is less than the probability that more than one cell would have been labeled with the same allele in the initial population, if the number of cells sampled is less than the initial population size. This means that $p_{singleton}(i;C)\big|_{C=M}$ underestimates the probability that observed cells share a unique progenitor and conversely $p_{duplicate}(i;C)\big|_{C=M}$ overestimates the probability that the observed cells arise from multiple progenitors. Therefore, in practice, if the initial population size $C$ is not known, computing $p_{singleton}(i;C)\big|_{C=M}$ using the number of sampled cells (in place of the unknown initial population size) is a conservative bound on the probability that allele $i$ is marking a unique clone, as long as $M < C$. In the case where $M > C$, the estimate is by definition more conservative because it assumes more cells were initially labeled than was actually the case.

2. The more non-uniform the proliferation rates of progenitor cells, the more conservative we are in using $p_{singleton}(i;C)\big|_{C=M}$ and $p_{duplicate}(i;C)\big|_{C=M}$ as bounds.

3. The fewer cells sampled at the final timepoint, the more conservative we are in using $p_{singleton}(i;C)\big|_{C=M}$ and $p_{duplicate}(i;C)\big|_{C=M}$ as bounds, achieving equality only in the limit of perfect sampling, where $M \gg C$.

*Statistical Significance of Alleles*

Previous barcoding systems fail to account for the natural ubiquity of alleles, mistakenly attributing kinship to cells coincidentally marked by the same allele. Since common alleles by definition occur in many cells, the number of cells implicated in these false-positive conclusions can be high. We circumvent this issue by assigning a p-value to each allele, which quantifies statistical significance of the allele according to the biological application, so that standard statistical best practices can be used.

Case 1: p-value for marking clones when $C$ is known or $C < M$

Consider the case where $C$, the number of cells initially labeled, is known (or there is an estimate on the upper bound $\leq M$, the number of cells observed at the sampling timepoint). If at the sampling timepoint, we observe at least one cell harboring allele $i$, $X_i > 0$, we would like to quantify the probability that the $X_i$ cells arose from more than one progenitor ($C_i > 1$) coincidentally marked with the same allele at the initial induction. If this probability is $< \alpha$, we can be confident at significance level $\alpha$, that all $X_i$ cells marked with allele $i$, originated from a single progenitor. This probability is just $p_{duplicate}$ introduced above, where we now use the approximation that the occurrence frequency of a given allele follows a Poisson distribution.

$$p_{duplicate}(i;C) = \mathrm{P}(C_i > 1 \mid X_i > 0; \lambda_i, C) = \mathrm{P}(C_i > 1 \mid C_i > 0; \lambda_i, C) = \frac{1 - e^{-\lambda_i C} - (\lambda_i C)e^{-\lambda_i C}}{1 - e^{-\lambda_i C}}$$

Note that conditional on allele $i$ being observed, $p_{duplicate}$ does not depend on the number of cells observed ($X_i$). Given the number of alleles and the distribution of their Poisson rates in the CARLIN system, for sufficiently large $C$, $p_{duplicate}$ will always be larger than a preset significance level for some of the alleles. For example, at $C = 5000$, 72% of alleles are significant at $\alpha = 0.05$, representing a cumulative allele frequency probability of 20% so that a fifth of cells will be marked uniquely at this significance level (Figure 3I).

Case 2: p-value for marking clones when $M < C$ or $C$ is unknown

In this case, we still seek to quantify the probability that $X_i$ cells arose from multiple progenitors, but without knowing the size of the initial pool. We assume that the $X_i$s form an unbiased sampling. The probability of observing more than one cell independently labeled with allele $i$ conditioned on having observed the allele is given by,

$$p_{clonal}(i;M) = P(X_i > 1 \mid X_i > 0; \lambda_i, M) = \frac{1 - e^{-\lambda_i M} - (\lambda_i M)e^{-\lambda_i M}}{1 - e^{-\lambda_i M}}$$

Note that $p_{clonal}$ is just $p_{duplicate}$ evaluated at $C = M$, according to our earlier justification (see *Limitations of a Finite Number of Alleles for Uniquely Marking Clones*) and constitutes an upper bound on the probability (a conservative p-value). We separately annotate this as $p_{clonal}$ instead of $p_{duplicate}$ to make clear that in this case, we have no knowledge of $C$ or $C_i$.

As $M$ grows, we are more likely to sample redundantly marked cells. Once $M \geq 35{,}000$, even the most rare allele is expected to mark multiple progenitors at significance level $\alpha = 0.05$. The upper limit on $M$ is determined by the smallest Poisson rate in our bank, the mean rate assigned to the unobserved alleles. Although there is actually a distribution of rates for unobserved alleles, we still cannot theoretically go beyond the limit imposed by the mean (as we cannot actually associate an unobserved allele to a particular rate in this distribution). This regime is indicated in grey in Figure 3I. We use this p-value for all our SC analysis where $M < 1500$.

Case 3: p-value for allele frequency

Lastly, we define a p-value that quantifies the probability of observing allele $i$ expressed in $X_i$ cells or more, under the null model that the allele has Poisson rate parameter $\lambda_i$.

$$p_{frequency}(i;M) = P(X \geq X_i \mid X_i > 0; \lambda_i, M) = \frac{1 - \sum_{k=0}^{X_i - 1} \frac{(\lambda_i M)^k e^{-\lambda_i M}}{k!}}{1 - e^{-\lambda_i M}}$$

If $p_{frequency}(i;M) < \alpha$, this means that cells marked with allele $i$ are significantly over-represented in a library of $M$ cells (at significance level $\alpha$). This does not necessarily indicate that all cells expressing the allele are related. However, it does indicate that at least one progenitor marked with the allele expanded more compared to progenitors marked with other alleles. The more skewed the proliferation rates ($b \to \infty$ according to notation introduced in *Limitations of a Finite Number of Alleles for Uniquely Marking Clones*), the more likely the over-representation is due to a single progenitor.

*Statistically Significant Clones for SC Analysis*

An allele was determined to be statistically significant if it had a $p_{clonal}$ that survived multiple hypothesis testing using the Benjamini-Hochberg procedure at a false discovery rate (FDR) of 0.05. A clone (a group of cells marked with the same allele) was said to be statistically significant, if its allele was statistically significant. For the 5-FU experiments, the number of observed cells ($M$) was set separately for each sample to equal the number of cells with an edited CARLIN allele in that sample (Supplementary Table 3). For the embryonic induction experiment, the number of observed cells ($M$) was the total number of cells with an edited CARLIN allele across the four bones (Supplementary Table 3).

*Comparing Progeny Origin in Development and Adult Hematopoiesis*

To quantify what fraction of statistically significant clones included both HSCs and non-HSCs, we computed the proportion of statistically significant alleles that marked cells belonging to both the HSC and non-HSC clusters. We also computed this proportion over the three controls as an average of the three proportions, weighted by the number of statistically significant alleles in each control. We performed a two-proportion z-test for the alternative hypothesis that the proportion is smaller in the control, rejecting at significance level $\alpha = 0.05$.

To quantify what fraction of the observed non-HSCs share an allele with an observed HSC (i.e. belong to an HSC-rooted clone) in the embryonic induction and 5-FU experiments, we computed the proportion of cells in non-HSC clusters that had a statistically significant allele which also appeared in a cell found in the HSC cluster. We also computed this proportion over the three controls as an average of the three proportions, weighted by the number of non-HSCs marked with statistically significant alleles in each control. We performed a two-proportion z-test for the alternative hypothesis that the proportion is larger in development/adult hematopoiesis than in the control, rejecting at significance level $\alpha = 0.05$.

*Comparing Number of Clones and Clone Size Distributions in Adult Hematopoiesis*

To determine if there were significantly fewer statistically significant clones in the 5-FU treated mouse, relative to the negative control, we first selected cells belonging to statistically significant clones. Let $M'$ be the minimum of the number of selected cells across the two treatments. We sampled $M'$ cells with replacement from each pool of selected cells, and counted the number of observed alleles. We repeated this procedure 10,000 times to get a distribution of the number of observed alleles in the two treatments. We performed a t-test for the alternative hypothesis that the mean of the number of observed alleles is less in the 5-FU treated sample at a significance level of $\alpha = 0.05$.

To determine if there are significant differences in the average HSC-rooted clone size between the 5-FU treated mouse and the negative control mouse, we selected cells marked by statistically significant alleles that also appear in at least one cell belonging to the HSC cluster. Let $M'$ be the minimum of the number of selected cells across the two treatments. We sample $M'$ cells with replacement from each pool of selected cells, letting $\{L_j\}_{j=1}^{M'}$ be the allele labels of the sampled cells, and $X_i = \sum_{j=1}^{M'} \delta_{L_i,L_j}$ the number of cells labeled with the same allele as cell $i$ (the size of the clone to which a cell belongs). We performed a one-sided Mann-Whitney test using $X_i$ for the alternative hypothesis that the clone sizes in the negative control are smaller at a significance level of $\alpha = 0.05$. We repeat this procedure 10,000 times to get a distribution of p-values, and report the 99$^{th}$ percentile value (see Supplementary Table 4).

To determine if the distribution of HSC-rooted clone sizes was significantly non-uniform, we selected cells marked by statistically significant alleles that also appear in at least one cell belonging to the HSC cluster. Suppose $M'$ cells are selected in a particular sample in this fashion, yielding $N \leq M'$ alleles. Let $L_j \in \{1 \dots N\}$, $1 \leq j \leq M'$, be the allele label of cell $j$. We sample $M'$ cells with replacement from this selected set of cells, and construct a CDF over the occurrence frequency of sampled allele labels $X_i = \sum_{j=1}^{M'} \delta_{L_j,i}$. We also sample allele labels $M'$ times from a discrete uniform distribution with $N$ bins, and again construct a CDF over the occurrence frequency of allele labels. We compute the p-value associated with the KS-statistic derived from comparing these two CDFs. We repeat this procedure 10,000 times, and report the 99$^{th}$ percentile for the p-value distribution.

*Statistical Significance of Bone/Fate Bias*

To quantify whether an allele $i$ displays significant fate bias, we consider the deviation of the observed number of cells marked with allele $i$, $(f_{i,1}, \dots f_{i,B})$, across $B$ possible bins (representing bones or phenotypes), against a multinomial null distribution given by $P(N_i = \sum_{j=1}^{B} f_{i,j}, p_1, \dots, p_B)$. For the embryonic induction dataset, we tested fate bias over different bones ($B = 4$). We also tested for bias over phenotypes, annotated from the Louvain clusters as defined in Figure 5B ($B = 5$; the lymphoid cluster which only had a few dozen cells was omitted), for each bone separately and pooling across bones.

The bin probabilities are the maximum likelihood estimators found by normalizing frequency counts after pooling all edited alleles, so that $p_j = \frac{\sum_i f_{i,j}}{\sum_j \sum_i f_{i,j}}$. The fate bias p-value that is reported is the probability under the null model of the range (the difference between the largest and smallest frequency across the bins for each allele) being greater or equal to the range observed. We compute this probability exactly according to the method described in (Corrado 2011). We note that while this method does not require that all bins have equal probability, using this measure to test bias is prone to false negatives when there is a large skew in bin probabilities, since the range does not take into account whether the maximum frequency occurred at a bin with low probability; in our data, all the bins (whether they are bones or coarse-grained phenotypes), have roughly the same probability. We only report fate bias p-values for statistically significant alleles (as defined in *Statistically Significant Clones for SC Analysis*) and reject at significance level $\alpha = 0.05$ (Bonferroni-corrected on the number of statistically significant alleles).

*Equivalence Testing for SC Amplicon Protocol Reproducibility*

To quantify the reproducibility of our single-cell CARLIN amplicon protocol, we prepared libraries in duplicate starting from the same single-cell transcriptome library for the samples in the 5-FU experiment. We performed equivalence testing on the frequency distribution of alleles (edited and unedited) called by the CARLIN pipeline, when applied independently on these replicates (Supplementary Figure 6A). We used the KS-statistic at a test value of 0.05 to determine whether two replicates are equivalent. We performed 10,000 bootstrapping trials, and computed the KS-statistic for each trial, to obtain a distribution over all trials. For all samples, the one-sided $(1\text{-}\alpha)$ confidence interval excluded the test value of 0.05, so replicates are said to be equivalent at significance level $\alpha = 0.05$. The results presented in the main text and Figure 6, were generated by pooling the sequencing data from the replicates, and running the CARLIN pipeline on the combined data.

**Lineage Reconstruction**

We pool together all cells/transcripts across all libraries/tissues, so that the reconstruction is agnostic to the origin of the alleles in the data. We use the specific details of mutations called in the alleles (see *Mutation Calling*) to establish a hierarchy across the alleles and thereby obtain a simplified phylogeny tree.

*Filtering Alleles*

For the in vivo data, we discarded alleles whose $p_{frequency}$ did not satisfy an FDR threshold of 0.05 under the Benjamini-Hochberg procedure. The number of observations, $M$, used in computing $p_{frequency}$, was the total number of edited transcripts after pooling the data. We used $p_{frequency}$ as opposed to $p_{clonal}$ because even if we are not confident that all cells that share a given allele came from the same progenitor, the relative expansion of one allele compared with another contains important information about clonal expansion across different tissues.

*Tree Reconstruction Algorithm*

Here, we describe a simple tree reconstruction algorithm used as a proof of concept for lineage tracing in the CARLIN system. Since a primary contribution of this paper is not in the domain of tree reconstruction, we defer investigation of how more sophisticated schemes like those newly developed in (Feng 2019) may improve the analysis.

For this algorithm, we make two assumptions, both of which are justified empirically by the data (see Figure 2D):

1) *Both leaf and internal nodes of the tree can only be occupied by alleles observed in the sample*. Since the efficiency of Cas9 is not 100%, we expect that every allele found after editing round $T$ will also be present after editing round $T+1$, because not every cell marked with a particular allele is further modified. The task of tree reconstruction is therefore simplified from inference of a novel set of mutations harbored by internal nodes, to identification of alleles which could likely be the internal nodes. In this scheme, we note that an allele must appear as a leaf node exactly once. Some leaf nodes may additionally appear once as an internal node with the same allele. In such cases, the internal node will always appear as a parent to the leaf node.

2) *A child allele may only differ from its parent allele by either having additional mutations to those harbored by its parent, and/or possessing a subsuming deletion, whose endpoints are located at pristine target sites in the parent*. The subsuming deletion would remove any history of parental mutations between deletion endpoints. This assumption therefore

requires that there are no repair mechanisms, precluding the possibility of mutations that are observed in the parent at sites that are unmodified in the child.

Assumption (2) allows us to determine whether any pair of alleles fall along the same lineage, in which one allele could be an ancestor of the other. Given a list of $n$ alleles (including the unmodified reference), we construct an $n$ x $n$ Boolean matrix $A$ where $A_{i,j} = 1$ if allele $j$ can be a descendent of allele $i$. We subsequently set to zero some of the non-zero entries using the following rules:

1) *Frequency Criteria:* We set $A_{i,j} = 0$ if the number of cells with allele $j$ is larger than the number of cells with allele $i$. We impose this criteria, consistent with Assumption (1), because we expect the number of cells with the parent allele to exceed the number of cells with the child allele for two reasons: (i) the cutting efficiency is typically <<100% so that the parent allele remains substantially represented among cells and (ii) multiple distinct outcomes can result from additional modifications to a starting allele, so that a single child allele is unlikely to become more prevalent than the parent, its prevalence being diluted by its siblings.

2) *Maximum Mutation Preservation*: For each child allele that can potentially be descended from multiple parent alleles, only the candidate parent alleles that share the maximum number of mutations with the child allele are retained as the potential parents, and the other entries are set to zero. This entails that the number of mutations required to transform from parent to child allele is the minimum among possible parents, based on the reasoning that multiple editing events leading to a child allele are more unlikely than a single editing event. This criterion ensures that a given allele is most likely to be connected to its nearest ancestor so that matrix $A$ can be interpreted as the set of parent-child relationships.

3) Alleles that have no parent allele after this pruning procedure are linked to the reference allele at the root of the tree.

A limitation of CARLIN is that alleles exhibiting large deletions could potentially be descended from almost any parent allele. In these cases, we have no choice but to connect the allele to the root of the tree as the lineage information in the allele is lost. However, such alleles are generally common in the bank and are therefore likely to be filtered out using our statistical significance tests (Figure 4B; *Filtering Alleles*).

Even after the pruning procedure, a given allele can still have multiple parent alleles. To construct a consensus tree from these potentially contradictory lineage relationships, we generate an ensemble of trees, by running simulations in which we probabilistically sample the parent edges of children with multiple candidate parents.

The tree is reconstructed recursively as follows: given a partially reconstructed tree and the current node, we randomly select one of the remaining unplaced alleles that is a child of the current node according to the $A$ matrix. We attach the selected allele to the current node as its child. The added node then becomes the current node. When none of the remaining unplaced alleles is a valid child of the current node, we backtrack by setting the current node to be the parent node. We initialize the algorithm by starting with the reference allele as the current node (defined as the root of the tree). Lastly, we include a leaf node for each internal node.

*Consensus Tree*

To determine lineage relationships that are conserved across our simulated trees, we focused on nodes that occur closer to the root, reasoning that they are less variable across different reconstructions and carry more information because they have more cells associated with their alleles (due to the frequency criteria).

The depth of a given node is one less than the number of nodes on the path from the root to the node. Define a depth-$X$ rooted path for a given internal node at depth-$X$, to be a sequence of alleles occurring on the path from the root of the tree (depth-0) to the internal node. We tabulate the number of times we observe a particular depth-$X$ rooted path over the ensemble of trees. In addition, for each simulation in which the depth-$X$ rooted path occurs, we tabulate the sum of the frequency of the alleles (referred to as the clade population) of all the leaves that descend from the terminating node of the path. Over the ensemble of trees, we obtain a distribution of clade populations for each depth-$X$ rooted path.

We consider a depth-$X$ rooted path to be stable if it appears in fraction $f$ of the simulated trees. For the reconstructions presented in the paper, we used $f = 1$, and constructed a consensus tree by aggregating all stable depth-$X$ rooted paths, for $X \leq 3$. We chose $f = 1$ by rank-ordering the occurrence frequency of the rooted paths of a given depth across all simulated trees (Supplementary Figure 4) and observing that (i) the top-ranked paths occurred across all simulations, (ii) the occurrence frequency decayed rapidly for paths that don't appear in all simulations, and (iii) the clade populations associated with the paths that did not occur in all simulations were not substantial.

Given our choice of $f = 1$, the same consensus tree could have been reconstructed by simply removing child alleles that had multiple potential parents in the $A$ matrix. In general, other choices of $f$ permit reconstruction of trees with contradictory lineage information.

Since there is no contradictory lineage information for $f = 1$, the consensus tree can be visualized unambiguously. For aesthetic reasons, in Figure 2D and 4D we omitted display of depth-1 rooted paths whose terminal nodes are leaves with a minimum clade population across simulations less than 100.

*Tissue Distance*

We use the same tissue distance metric outlined in (Feng 2019). In short, for the consensus tree, we compute the distance between tissues $i$ and $j$ by first identifying all leaves corresponding to alleles found in tissue $i$. For each allele, we compute the distance to its closet ancestor which has a progeny allele that was found in tissue $j$ (one less than the number of nodes along the path connecting the leaf to the internal node). Next, we average this distance over all alleles for tissue $i$. We weight each term in the average by the product of the number of cells found in tissue $i$ marked by the leaf allele and the number of cells found in tissue $j$ marked by progeny alleles of the closest ancestor. In Figure 4H we show the matrix obtained from this procedure after symmetrization.

**Transcriptomic Analysis and Visualization**

After discarding cells with few UMIs and high expression of mitochondrial genes (see *Preprocessing*), the resulting gene expression matrices were processed using Seurat (v3.0.2 with default parameters except where indicated) as follows. First, cell cycle scores were assigned to each cell using the CellCycleScoring function. Next, UMI counts were normalized according to a regularized negative binomial regression model while regressing out the effect of cell cycle and mitochondrial genes (using the SCTransform function with vars.to.regress = c("percent.mt", "S.Score", "G2M.Score")).

The 4 bones of the embryonic induction dataset were aligned (and similarly the 5-FU and control samples), by finding 2000 common features using the FindIntegrationAnchors function, and specifying these, together with marker genes (Supplementary Figure 5E,6D), as landmarks to the IntegrateData function (Supplementary Figure 5C,6B). Joint dimensionality reduction was performed on the aligned datasets using RunPCA. Points in this embedding were used to construct UMAP plots (RunUMAP), and find neighbors for clustering (FindNeighbors). For all relevant functions, 20 principal components were used.

FindClusters (algorithm=2, resolution=1.0) was used to run the Louvain clustering algorithm with multilevel refinement (Supplementary Figures 5D, 6C). The dot plots in Supplementary Figures 5E and 6D were generated using data from Seurat's DotPlot function. Subclustering was performed on the 5-FU dataset by re-rerunning FindClusters with resolution=1.1 (Supplementary Figure 6E). Differential gene expression was performed using the FindMarkers function, whose output when applied to our data is shown in Supplementary Tables 5 and 6. Visualization of differentially expressed genes was handled using Seurat's VlnPlot function for Figure 6G and Supplementary Figure 6F,G, and DoHeatMap function for Figure 6H.

**Data and Code Availability**

All data used in this paper will be made available on the NCBI GEO upon publication. The CARLIN software package, together with the allele bank, is available at https://gitlab.com/hormozlab/carlin. All code and instructions to reproduce numerics and figures in the manuscript will be made available upon publication.

Smith, Tom Sean, Andreas Heger, and Ian Sudbery. 2017. "UMI-tools: Modelling sequencing errors in Unique Molecular Identifiers to improve quantification accuracy." *Genome research* gr-209601.

Zhang, Jiajie, Kassian Kobert, Tomáš Flouri, and Alexandros Stamatakis. 2013. "PEAR: a fast and accurate Illumina Paired-End reAd mergeR." *Bioinformatics* 30 (5): 614-620.

Zilionis, R., Nainys, J., Veres, A., Savova, V., Zemmour, D., Klein, A.M., and Mazutis, L. 2017. "Single-cell barcoding and sequencing using droplet microfluidics." *Nat Protoc* 12: 44-73.

### SUPPLEMENTARY FIGURES

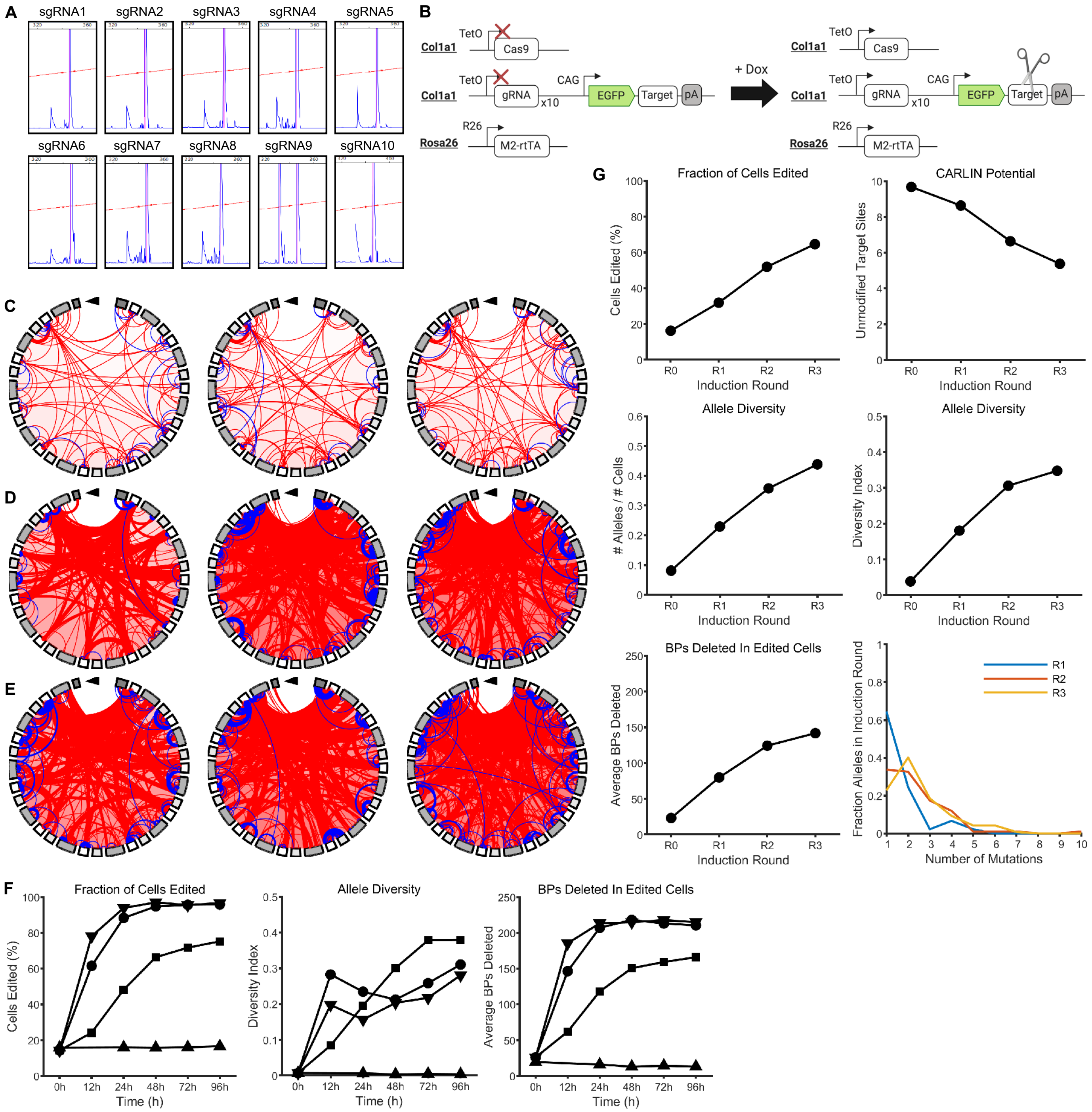

**Supplementary Figure 1: Additional quantification of editing in ES cells**

**A.** Fragment analysis results indicating cutting efficiency of individual sgRNAs.

**B.** Schematic of inducible CARLIN system.

**C.** Chord plots of CARLIN alleles at 24h, 48h, and 72h (left to right) in the absence of doxycycline (Dox), as in Figure 1F.

**D.** Chord plots of CARLIN alleles at 12h, 24h, and 48h (left to right) after induction with 0.20μg/mL (medium) of Dox, as in Figure 1F.

**E.** Chord plots of CARLIN alleles at 12h, 24h, and 48h (left to right) after induction with 1.00μg/mL (high) of Dox, as in Figure 1F.

**F.** From left to right, fraction of cells edited, diversity index (see Methods), and average base pairs deleted from CARLIN alleles across edited cells, for chronic induction experiments shown in Figure 1F-H. Legend as in Figure 1G-H

**G.** Fraction of cells edited (top left), CARLIN potential (top right), diversity metrics (middle row), average base pairs deleted in CARLIN allele across edited cells (bottom left), and frequency distribution of mutation counts (bottom right) for the pulsed induction experiment shown in Figure 2A.

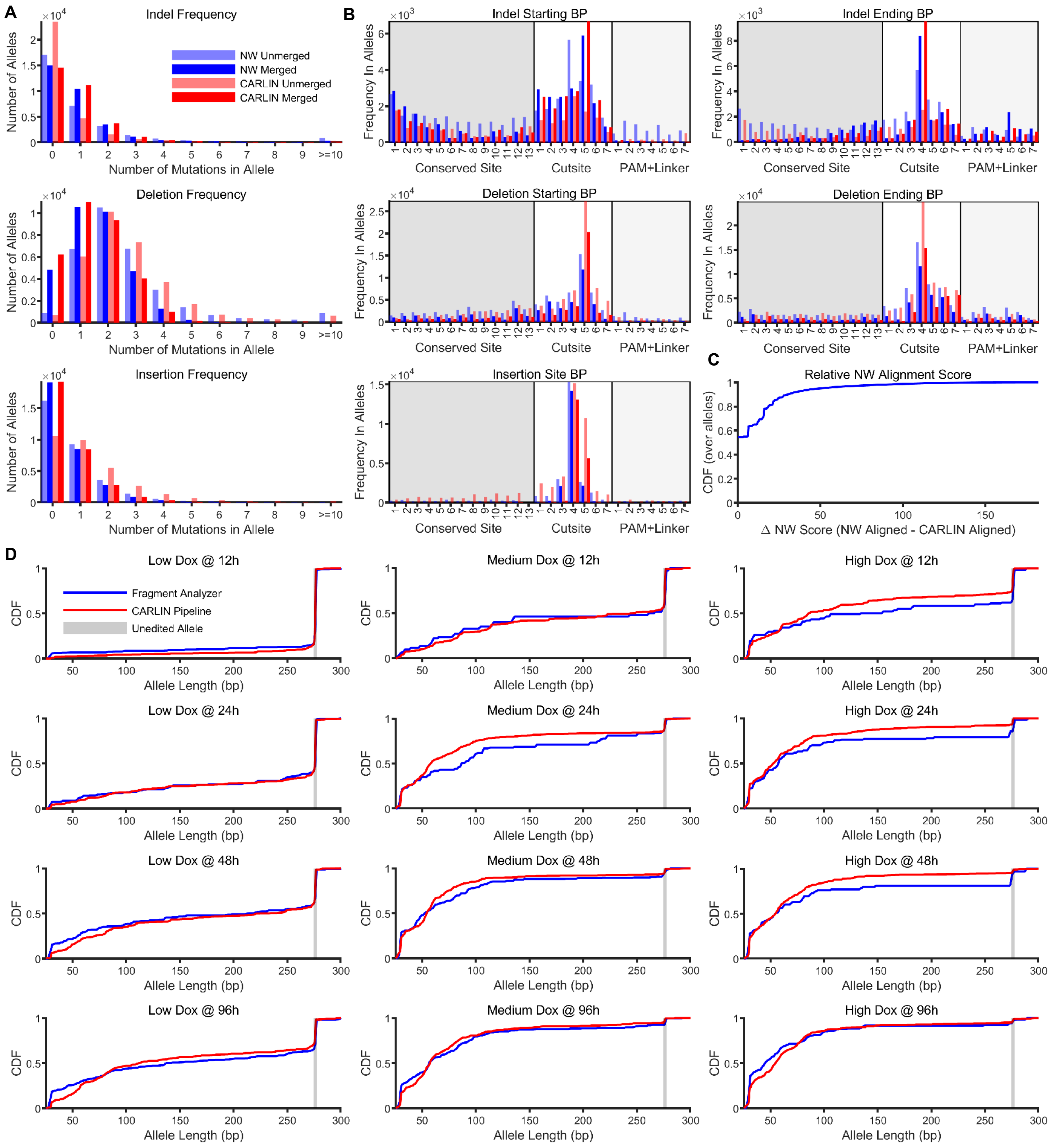

A
Indel Frequency
Number of Alleles
Number of Mutations in Allele
NW Unmerged
NW Merged
CARLIN Unmerged
CARLIN Merged
Deletion Frequency
Insertion Frequency
B
Indel Starting BP
Indel Ending BP
Deletion Starting BP
Deletion Ending BP
Insertion Site BP
Frequency In Alleles
Conserved Site
Cutsite
PAM+Linker
C
Relative NW Alignment Score
CDF (over alleles)
Δ NW Score (NW Aligned - CARLIN Aligned)
D
Low Dox @ 12h
Medium Dox @ 12h
High Dox @ 12h
Low Dox @ 24h
Medium Dox @ 24h
High Dox @ 24h
Low Dox @ 48h
Medium Dox @ 48h
High Dox @ 48h
Low Dox @ 96h
Medium Dox @ 96h
High Dox @ 96h
Fragment Analyzer
CARLIN Pipeline
Unedited Allele
CDF
Allele Length (bp)

**Supplementary Figure 2: Comparison of CARLIN analysis pipeline with NW alignment and fragment analysis**

**A.** Frequency of indels, deletions and insertions called using the standard Needleman-Wunsch (NW) and the CARLIN algorithm, when applied to the CARLIN sequences of alleles in the bank. For the series labelled 'Unmerged' we refrained from merging mutations (Methods).

**B.** Frequency of indels, deletions and insertions called as a function of their location along different motifs on the CARLIN array. Repetitions of the same motif are overlaid. Relative to the NW algorithm, the CARLIN algorithm tends to cluster mutations more frequently at the expected cutsites. Shading as in Figure 1B.

**C.** CDF of the difference between the NW score of the optimal alignment produced by the NW algorithm and the optimal alignment produced by the CARLIN algorithm. About half the alleles in the bank have alignments that match the NW algorithm. The remainder would have been discarded as suboptimal by the NW algorithm.

**D.** Comparison of fragment lengths as measured by fragment analysis (Methods), and the frequency-weighted distribution of allele lengths called by the CARLIN pipeline applied to NGS data. The data correspond to four timepoints from the three +Dox concentrations used in the chronic induction experiment shown in Figure 1F-H.

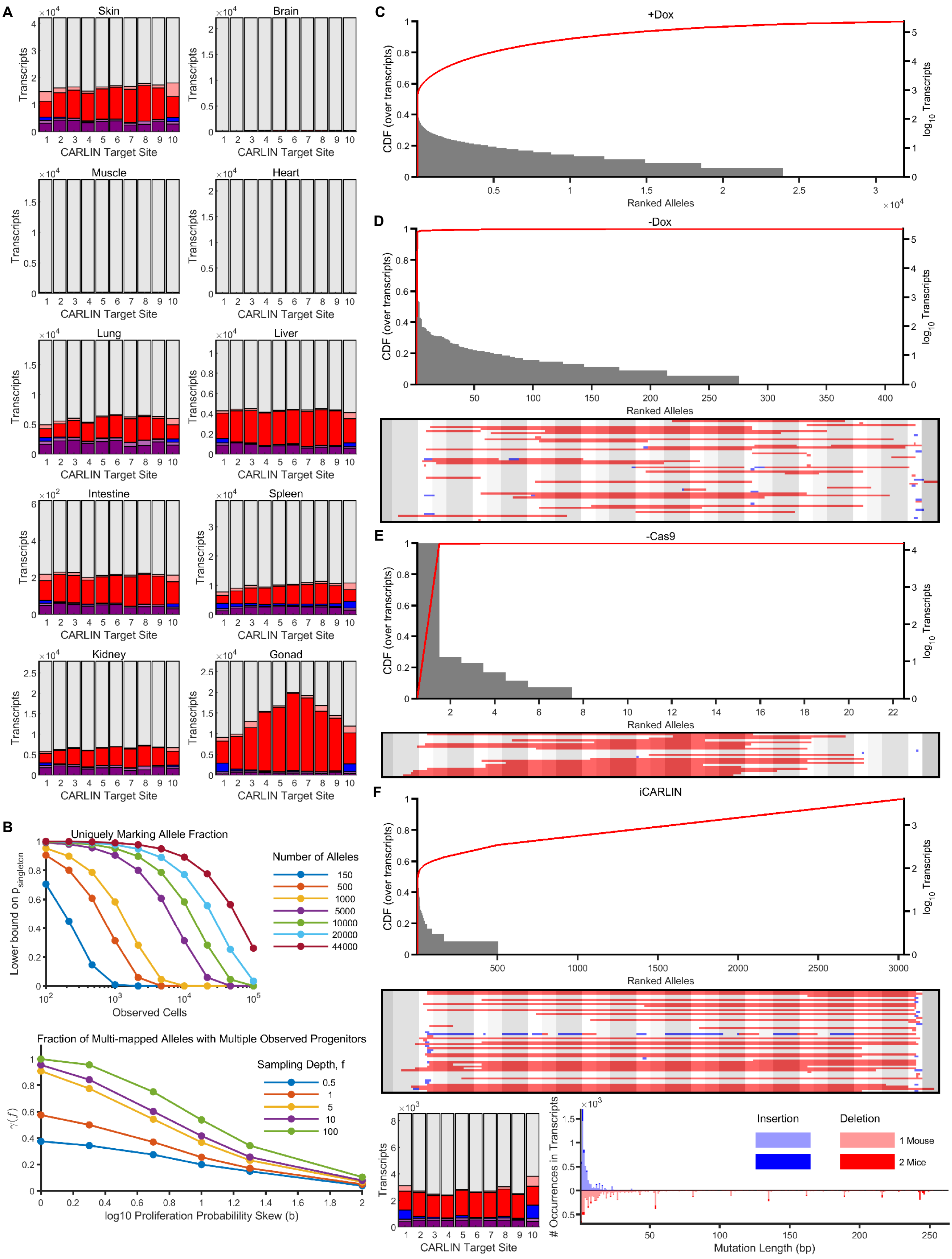
A
Skin
Brain
Muscle
Heart
Lung
Liver
Intestine
Spleen
Kidney
Gonad
Transcripts
CARLIN Target Site
B
Uniquely Marking Allele Fraction
Lower bound on p singleton
Observed Cells
Number of Alleles
150
500
1000
5000
10000
20000
44000
Fraction of Multi-mapped Alleles with Multiple Observed Progenitors
γ(f)
log10 Proliferation Probabililty Skew (b)
Sampling Depth, f
0.5
1
5
10
100
C
+Dox
CDF (over transcripts)
log10 Transcripts
Ranked Alleles
D
-Dox
E
-Cas9
F
iCARLIN
### Occurrences in Transcripts
Mutation Length (bp)
Insertion
Deletion
1 Mouse
2 Mice

**Supplementary Figure 3: Mutations in different tissues, and additional details of the allele bank**

**A.** Site decomposition plot as in Figure 3D for the tissues shown in Figure 3B, under doxycycline (Dox) induction. Figure legend same as in Figure 3D, which shows results for granulocytes comprising the bank.

**B.** (Top) Lower bound on the probability of observing an allele marking cells belonging to a unique progenitor ($p_{singleton}$, see Methods) for different numbers of observed cells, in the case of equally likely alleles. As the number of cells observed increases, the chance of observing cells that coincidentally share the same allele increases. This effect is counteracted by increasing the number of alleles available (different colored traces). (Bottom) When multiple progenitors are marked with the same allele, the more uneven the proliferation rates (larger values of the $b$ parameter in the proliferation model, see Methods), the less likely it is to observe the same allele shared across multiple cells that arise from different progenitors (see Methods for definition of $\gamma(f)$). However, as the number of observed cells increases (increasing $f$, see Methods), even progeny from slowly-proliferating progenitors that were redundantly marked are more likely to be observed, so that the fraction of alleles that ambiguously mark the ancestry of observed cells increases.

**C.** Distribution of observed allele frequencies for the bank. The top-ranked allele is the unedited reference.

**D.** (Top) Distribution of allele frequencies as in (C) and (bottom) the 50 most common edited alleles for a –Dox control obtained by harvesting ~310K granulocytes from each of two uninduced mice.

**E.** (Top) Distribution of allele frequencies as in (C) and the 50 most common edited alleles (bottom) for a –Cas9 control obtained by harvesting ~125K granulocytes from a single mouse.

**F.** Distribution of allele frequencies (top), the 50 most common edited alleles (middle), site decomposition plot as in Figure 3D (bottom left), and histogram of mutation frequencies (bottom right), obtained from harvesting granulocytes from two Dox-induced iCARLIN mice (Methods).

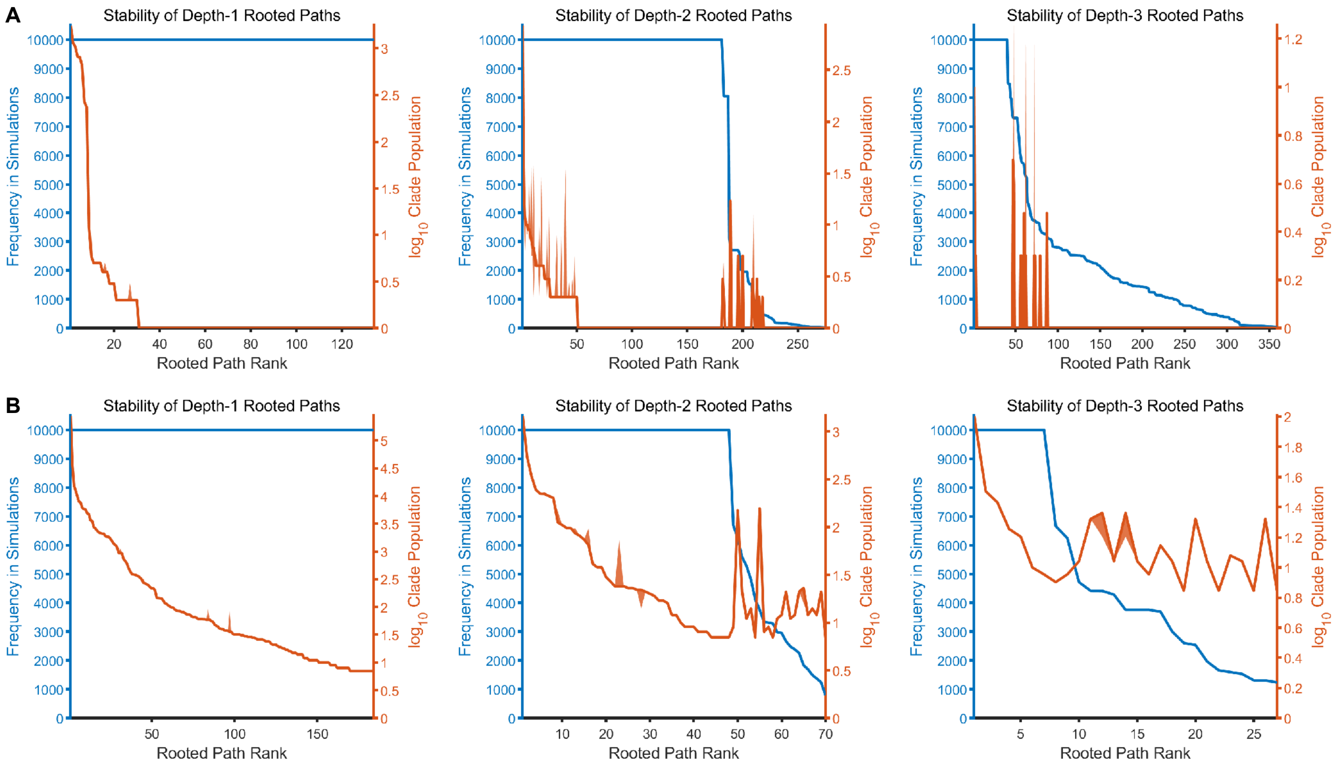


**Supplementary Figure 4: Additional details of lineage tree reconstruction simulations**

**A.** Number of instances (blue) of rooted paths (Methods) of depth 1-3 (left to right), found across 10,000 simulations for the *in vitro* tree reconstruction shown in Figure 2D. We deem a rooted path stable, if it is found in all 10,000 simulations. In orange, we show the median clade population (across all simulations in which it appears) for each rooted path (Methods). Error bars (shaded) are min/max across the simulations. **B.** Same as in (A) but for the in vivo tree reconstruction shown in Figure 4D.

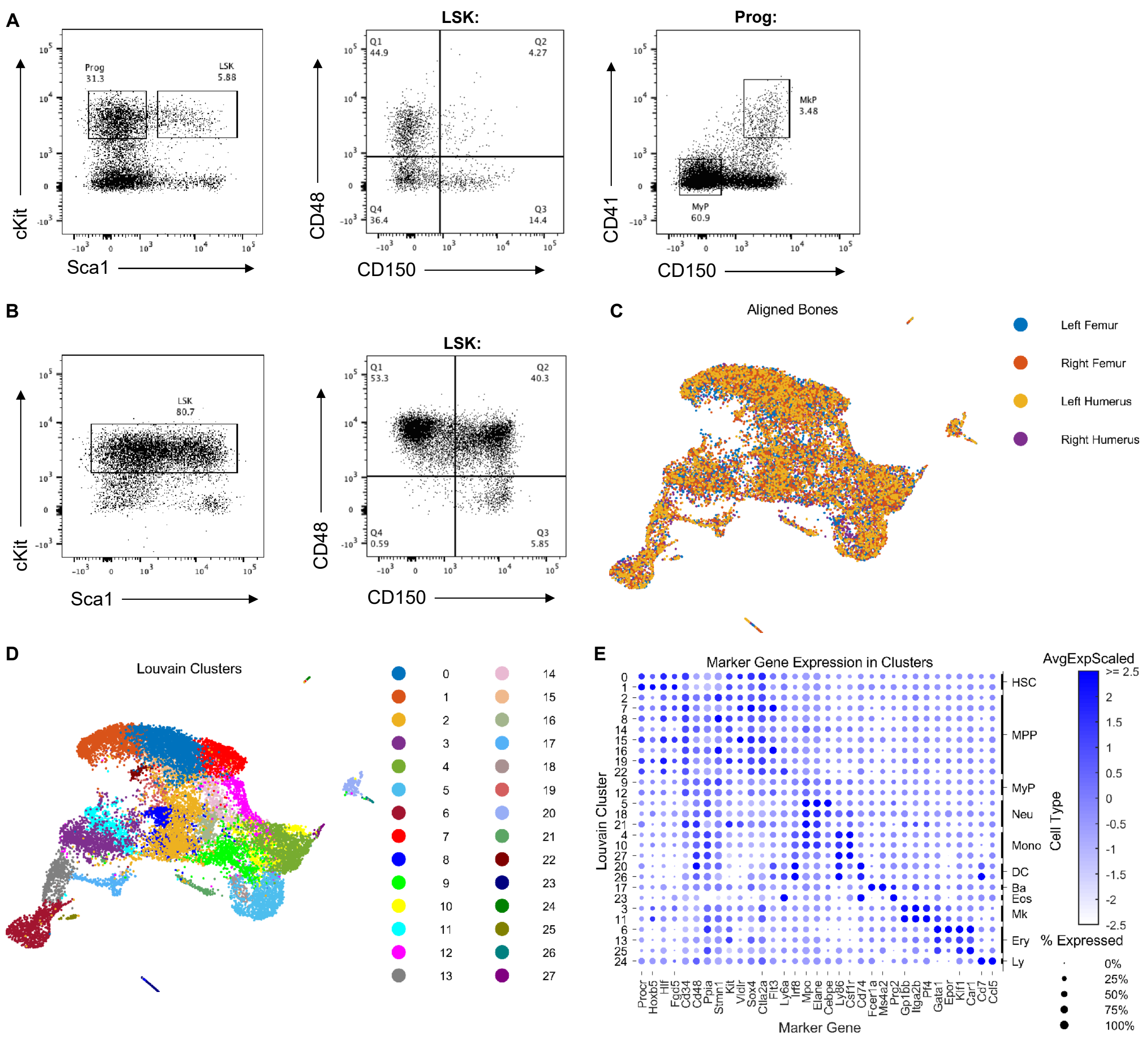


**Supplementary Figure 5: FACS plots and extended data for embryonic induction experiment**

**A.** FACS strategy used for sorting cells for encapsulation from bone marrow (BM) in the embryonic induction experiment (Figure 5) and control 5-FU experiment (Figure 6).

**B.** FACS strategy used for sorting cells for encapsulation from BM in the 5-FU experiment (Figure 6).

**C.** Overlay of transcriptome data from the 4 separate bones following batch correction.

**D.** Unsupervised Louvain clustering on the aligned transcriptomes from the 4 bones used in the embryonic induction experiment, generated 28 distinct clusters.

**E.** Dot plots of canonical blood cell-type specific markers used for fine-grain cell annotations of each Louvain cluster (HSC, hematopoietic stem cell; MPP, multipotent progenitor cell; MyP, myeloid progenitor; Neu, neutrophil; Mono, monocyte; DC, dendritic cell; Ba, basophil; Eos, eosinophil; Mk, megakaryocyte; Ery, erythrocyte; Ly, lymphoid cell). MyP, Neu, Mono, DC, Ba and Eos were further grouped to form the coarse-grained clusters shown in Figure 5B.

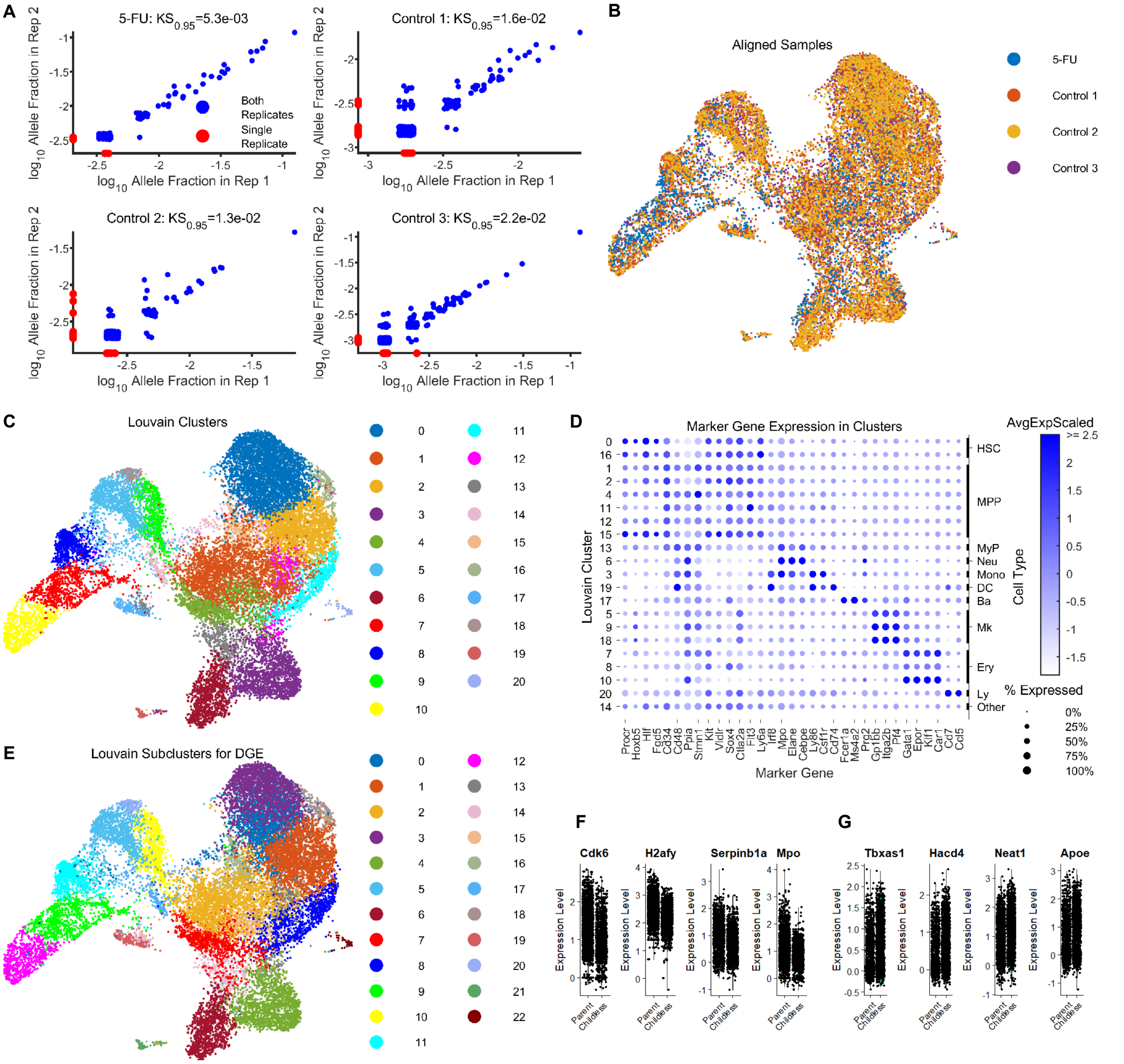

A
5-FU: KS0.95=5.3e-03
Control 1: KS0.95=1.6e-02
Control 2: KS0.95=1.3e-02
Control 3: KS0.95=2.2e-02
log10 Allele Fraction in Rep 1
log10 Allele Fraction in Rep 2
Both Replicates
Single Replicate
B
Aligned Samples
5-FU
Control 1
Control 2
Control 3
C
Louvain Clusters
D
Marker Gene Expression in Clusters
Louvain Cluster
Marker Gene
Cell Type
HSC
MPP
MyP
Neu
Mono
DC
Ba
Mk
Ery
Ly
Other
AvgExpScaled
% Expressed
Procr
Hoxb5
Hlf
Fgd5
Cd34
Cd48
Ppia
Stmn1
Kit
Vldr
Sox4
Clla2a
Flt3
Ly6a
Irf8
Mpo
Elane
Cebpe
Ly86
Csf1r
Cd74
Fcer1a
Ms4a2
Prg2
Gp1bb
Itga2b
Pf4
Gata1
Epor
Klf1
Car1
Cd7
Ccl5
E
Louvain Subclusters for DGE
F
Cdk6
H2afy
Serpinb1a
Mpo
G
Tbxas1
Hacd4
Neat1
Apoe
Expression Level
Parent
Childless

**Supplementary Figure 6: Extended data for 5-FU experiment**

**A.** The libraries for the 5-FU experiment (Figure 6) were prepared in replicate to analyze reproducibility. Each point corresponds to the proportion of cells in each replicate in which a given allele is found. For visualization purposes, the point corresponding to the unedited reference allele has been omitted and jitter has been added to distinguish stacked points.

**B.** Overlay of three control and one 5-FU treated single cell experiments following batch correction (Methods).

**C.** Unsupervised Louvain clustering after aligning the transcriptomes of the 5FU and control samples generated 21 distinct clusters.

**D.** Dot plots of canonical blood cell-type specific markers used for cluster annotation in 5-FU and control mice, using the same two-tier scheme described in Supplementary Figure 5E.

**E.** Louvain clustering after increasing the resolution parameter of the Louvain clustering algorithm (Methods). The HSC cluster 0 obtained at resolution 1.0 (see C), resolved into two clusters, 0 and 3, at resolution 1.1. We designated cluster 0 as the 'Parent Cluster', and cluster 3 as the 'Childless Cluster'. The other clusters did not change substantially.

**F.** Violin plots of the genes most differentially overexpressed in the HSC Parent cluster relative to the HSC Childless cluster, not shown in Figure 6G.

**G.** Violin plots of the genes most differentially underexpressed in the HSC Parent cluster relative to the HSC Childless cluster, not shown in Figure 6G.

#### SUPPLEMENTARY TABLES

##### Supplementary Table 1: Related to CARLIN construct design (annotated 5' to 3')

| | |
|---|---|
| **inducible sgRNAs** | |
| isgRNA1_F | tcccgactgcacgacagtcgacga |
| isgRNA1_R | aaactcgtcgactgtcgtgcagtc |
| isgRNA2_F | tcccgacacgactcgcgcatacga |
| isgRNA2_R | aaactcgtatgcgcgagtcgtgtc |
| isgRNA3_F | tcccgactacagtcgctacgacga |
| isgRNA3_R | aaactcgtcgtagcgactgtagtc |
| isgRNA4_F | tcccgcgagcgctatgagcgacta |
| isgRNA4_R | aaactagtcgctcatagcgctcgc |
| isgRNA5_F | tcccgatacgatacgcgcacgcta |
| isgRNA5_R | aaactagcgtgcgcgtatcgtatc |
| isgRNA6_F | tcccgagagcgcgctcgtcgacta |
| isgRNA6_R | aaactagtcgacgagcgcgctctc |
| isgRNA7_F | tcccgcgactgtacgcacacgcga |
| isgRNA7_R | aaactcgcgtgtgcgtacagtcgc |
| isgRNA8_F | tcccgatagtatgcgtacacgcga |
| isgRNA8_R | aaactcgcgtgtacgcatactatc |
| isgRNA9_F | tcccgagtcgagacgctgacgata |
| isgRNA9_R | aaactatcgtcagcgtctcgactc |
| isgRNA10_F | tcccgatacgtagcacgcagacga |
| isgRNA10_R | aaactcgtctgcgtgctacgtatc |
| **constitutive sgRNAs** | |
| csgRNA1_F | caccgactgcacgacagtcgacga |
| csgRNA1_R | aaactcgtcgactgtcgtgcagtc |
| csgRNA2_F | caccgacacgactcgcgcatacga |
| csgRNA2_R | aaactcgtatgcgcgagtcgtgtc |
| csgRNA3_F | caccgactacagtcgctacgacga |
| csgRNA3_R | aaactcgtcgtagcgactgtagtc |
| csgRNA4_F | caccgcgagcgctatgagcgacta |
| csgRNA4_R | aaactagtcgctcatagcgctcgc |
| csgRNA5_F | caccgatacgatacgcgcacgcta |
| csgRNA5_R | aaactagcgtgcgcgtatcgtatc |
| csgRNA6_F | caccgagagcgcgctcgtcgacta |
| csgRNA6_R | aaactagtcgacgagcgcgctctc |
| csgRNA7_F | caccgcgactgtacgcacacgcga |
| csgRNA7_R | aaactcgcgtgtgcgtacagtcgc |
| csgRNA8_F | caccgatagtatgcgtacacgcga |

| | |
|---|---|
| csgRNA8_R | aaactcgcgtgtacgcatactatc |
| csgRNA9_F | caccgagtcgagacgctgacgata |
| csgRNA9_R | aaactatcgtcagcgtctcgactc |
| csgRNA10_F | caccgatacgtagcacgcagacga |
| csgRNA10_R | aaactcgtctgcgtgctacgtatc |
| **Assembly Golden Gate isgRNA multiplex** | |
| isgRNA1_F | atatGgtctcagagaggtaccgcatgagattatcaaaaagg |
| isgRNA1_R | atatggtctctgtacgtacgtacaaaaaagcaccgactcggtg |
| isgRNA2_F | atatGgtctcagtacgcatgagattatcaaaaagg |
| isgRNA2_R | atatggtctctcaggcaggcaggaaaaaagcaccgactcggtg |
| isgRNA3_F | atatGgtctcacctggcatgagattatcaaaaagg |
| isgRNA3_R | atatggtctctgaccgaccgaccaaaaaagcaccgactcggtg |
| isgRNA4_F | atatGgtctcaggtcgcatgagattatcaaaaagg |
| isgRNA4_R | atatggtctctttgtttgtttgtaaaaaagcaccgactcggtg |
| isgRNA5_F | atatGgtctcaacaagcatgagattatcaaaaagg |
| isgRNA5_R | atatggtctctacgtacgtacgtaaaaaagcaccgactcggtg |
| isgRNA6_F | atatGgtctcaacgtgcatgagattatcaaaaagg |
| isgRNA6_R | atatggtctcttggctggctggcaaaaaagcaccgactcggtg |
| isgRNA7_F | atatGgtctcagccagcatgagattatcaaaaagg |
| isgRNA7_R | atatggtctctaaagaaagaaagaaaaaagcaccgactcggtg |
| isgRNA8_F | atatGgtctcactttgcatgagattatcaaaaagg |
| isgRNA8_R | atatggtctctagctagctagctaaaaaagcaccgactcggtg |
| isgRNA9_F | atatGgtctcaagctgcatgagattatcaaaaagg |
| isgRNA9_R | atatggtctctctcgctcgctcgaaaaaagcaccgactcggtg |
| isgRNA10_F | atatGgtctcacgaggcatgagattatcaaaaagg |
| isgRNA10_R | atatggtctctgcggttaattaaataaaaaagcaccgactcggtg |
| **Assembly Golden Gate csgRNA multiplex** | |
| csgRNA1_F | atatGgtctcagagaggtaccTTTCCCATGATTCCTTCATA |
| csgRNA1_R | atatggtctctgtacgtacgtacAACGGGTACCTCTAGAGCCA |
| csgRNA2_F | atatGgtctcagtacTTTCCCATGATTCCTTCATA |
| csgRNA2_R | atatggtctctcaggcaggcaggAACGGGTACCTCTAGAGCCA |
| csgRNA3_F | atatGgtctcacctgTTTCCCATGATTCCTTCATA |
| csgRNA3_R | atatggtctctgaccgaccgaccAACGGGTACCTCTAGAGCCA |
| csgRNA4_F | atatGgtctcaggtcTTTCCCATGATTCCTTCATA |
| csgRNA4_R | atatggtctctttgtttgtttgtAACGGGTACCTCTAGAGCCA |
| csgRNA5_F | atatGgtctcaacaaTTTCCCATGATTCCTTCATA |
| csgRNA5_R | atatggtctctacgtacgtacgtAACGGGTACCTCTAGAGCCA |
| csgRNA6_F | atatGgtctcaacgtTTTCCCATGATTCCTTCATA |
| csgRNA6_R | atatggtctcttggctggctggcAACGGGTACCTCTAGAGCCA |
| csgRNA7_F | atatGgtctcagccaTTTCCCATGATTCCTTCATA |

| | |
|---|---|
| csgRNA7_R | atatggtctctaaagaaagaaagAACGGGTACCTCTAGAGCCA |
| csgRNA8_F | atatGgtctcactttTTTCCCATGATTCCTTCATA |
| csgRNA8_R | atatggtctctagctagctagctAACGGGTACCTCTAGAGCCA |
| csgRNA9_F | atatGgtctcaagctTTTCCCATGATTCCTTCATA |
| csgRNA9_R | atatggtctctctcgctcgctcgAACGGGTACCTCTAGAGCCA |
| csgRNA10_F | atatGgtctcacgagTTTCCCATGATTCCTTCATA |
| csgRNA10_R | atatggtctctgcggttaattaaatAACGGGTACCTCTAGAGCCA |
| **CARLIN reference sequence** | |
| cgccggactgcacgacagtcgacgatggagtcgacacgactcgcgcatacgatggagtcgactacagtcgctacgacgatggagtcgcgag cgctatgagcgactatggagtcgatacgatacgcgcacgctatggagtcgagagcgcgctcgtcgactatggagtcgcgactgtacgcacacg cgatggagtcgatagtatgcgtacacgcgatggagtcgagtcgagacgctgacgatatggagtcgatacgtagcacgcagacgatgggagct | |
| **Pre-polyA UTR sequence** | |
| agaattctaactagagctcgctgatcagcct | |
| **pBS31-inducible CARLIN plasmid** | |
| https://benchling.com/s/lZx1S0ul | |
| **pBS31-constitutive CARLIN plasmid** | |
| https://benchling.com/s/zs5pUinp | |
| **CARLIN amplification** | |
| CARLIN_fwd1 | gtcctgctggagttcgtgac |
| CARLIN_fwd2 | gagctgtacaagtaagcggc |
| CARLIN_rev | gcaactagaaggcacagtcg |
| **Fragment analysis** | |
| FAM_CARLIN_fwd | /56-FAM/gagctgtacaagtaagcggc |
| **Bulk DNA** | |
| NGS_UIM_D_F | CTACACGACGCTCTTCCGATCTNNNNNNNNNNgagctgtacaagtaagcggc |
| NGS2_F | AATGATACGGCGACCACCGAGATCTACACTCTTTCCCTACACGACGCTCT TCCGATCT |
| NGS1_R | GTGACTGGAGTTCAGACGTGTGCTCTTCCGATCTgcaactagaaggcacagtcg |
| P5 | AATGATACGGCGACCACCGA |
| NG2R_I# | CAAGCAGAAGACGGCATACGAGATXXXXXXGTGACTGGAGTTC<br><br>XXXXXX = TruSeq Illumina indices |
| **Bulk RNA** | |
| RT_Bulk_CARLIN | CTACACGACGCTCTTCCGATCTNNNNNNNNNNGCAACTAGAAGGCACAG TCG |
| NGS1_F | gtcctgctggagttcgtgac |
| NGS2_F | AATGATACGGCGACCACCGAGATCTACACTCTTTCCCTACACGACGCTCT TCCGATCT |
| NGS1_Bulk_R | GTGACTGGAGTTCAGACGTGTGCTCTTCCGATCTgagctgtacaagtaagcggc |
| P5 | AATGATACGGCGACCACCGA |
| NG2R_I# | CAAGCAGAAGACGGCATACGAGATXXXXXXGTGACTGGAGTTC |

| | |
|---|---|
| | XXXXXX = TruSeq Illumina indices |
| **InDrop v3** | |
| R1-N6 | TCGTCGGCAGCGTCAGATGTGTATAAGAGACAGNNNNNN |
| R1-CARLIN | TCGTCGGCAGCGTCAGATGTGTATAAGAGACAGgagctgtacaagtaagcggc |
| R2 | CAAGCAGAAGACGGCATACGAGATGGGTGTCGGGTGCAG |
| P7 | CAAGCAGAAGACGGCATACGAGAT |
| R1_I# | AATGATACGGCGACCACCGAGATCTACACXXXXXXXXTCGTCGGCAGCGTC<br><br>XXXXXXXX = InDrop v3 indices |
| **10X Chromium** | |
| 10X-CARLIN_1-bio | bio-gtcctgctggagttcgtgac |
| 10X-CARLIN_2 | GTGACTGGAGTTCAGACGTGTGCTCTTCCGATCTgagctgtacaagtaagcggc |
| P5-PR1 | AATGATACGGCGACCACCGAGATCTACACTCTTTCCCTACACGACGCTC |
| P5 | AATGATACGGCGACCACCGA |
| **Genotyping Col1a1 locus** | |
| Col1a1_reverse | ccctccatgtgtgaccaagg |
| Col1a1_fwd_WT | gcacagcattgcggacatgc |
| Col1a1_fwd_mut | gcagaagcgcggccgtctgg |
| **Genotyping Rosa26 locus** | |
| Rosa26_fwd | aaagtcgctctgagttgttat |
| Rosa26_rev_WT | ggagcgggagaaatggatatg |
| Rosa26_rev_mut | gcgaagagtttgtcctcaacc |

**Supplementary Table 2: Penalties ($P_B/P_E$) used in alignment algorithm (Supplementary Figure 2 & Methods)**

| Motif | Base Pair | | | | | | | | | | | | |
|---|---|---|---|---|---|---|---|---|---|---|---|---|---|
| | *1* | *2* | *3* | *4* | *5* | *6* | *7* | *8* | *9* | *10* | *11* | *12* | *13* |
| **Opening Insertion** | 10 | | | | | | | | | | | | |
| **Prefix** | 10 | 10 | 10 | 10 | 10 | | | | | | | | |
| **Conserved Site** | 10 | 9.875 | 9.75 | 9.625 | 9.5 | 9.375 | 9.25 | 9.125 | 9 | 8.875 | 8.75 | 8.625 | 8.5 |
| **Cutsite** | 8.0 | 7.5 | 7.0 | 6.5 | 6.0/<br>6.5 | 6.0/<br>6.5 | 6.0/<br>6.5 | | | | | | |
| **PAM+Linker** | 8.6 | 8.8 | 9.0 | 9.2 | 9.4 | 9.6 | 9.8 | | | | | | |
| **Postfix** | 8.6 | 8.8 | 9.0 | 9.2 | 9.4 | 9.6 | 9.8 | 10 | | | | | |

**Supplementary Table 3: Metrics for single cell data (Figures 5 & 6)**

<table>
<tr><th rowspan="2">Sample</th><th rowspan="2">System</th><th colspan="4">Transcriptome</th><th colspan="5">CARLIN Amplicon</th></tr>
<tr><th>Cells Loaded</th><th>Minimum UMI Count</th><th>Filtered CBs</th><th>Median Genes in Filtered CBs</th><th>Filtered CBs</th><th>Mean Reads in Filtered CBs</th><th colspan="2">Alleles</th><th>% CBs Edited</th></tr>
<tr><td>Ctrl 1</td><td>10X v3</td><td>8000</td><td>1632</td><td>5960</td><td>2938</td><td>3765</td><td>330</td><td colspan="2">532</td><td>19</td></tr>
<tr><td>Ctrl 2</td><td>10X v3</td><td>8000</td><td>2113</td><td>6025</td><td>2813</td><td>3039</td><td>363</td><td colspan="2">395</td><td>17</td></tr>
<tr><td>Ctrl 3</td><td>10X v3</td><td>8000</td><td>2012</td><td>4832</td><td>3305</td><td>1539</td><td>1329</td><td colspan="2">693</td><td>74</td></tr>
<tr><td>5-FU 1</td><td>10X v3</td><td>8000</td><td>1539</td><td>4073</td><td>3330</td><td>2000</td><td>621</td><td colspan="2">89</td><td>16</td></tr>
<tr><td>5-FU 2</td><td>10X v2</td><td>8000</td><td>3267</td><td>4148</td><td>2518</td><td>519</td><td>2471</td><td colspan="2">33</td><td>18</td></tr>
<tr><td>5-FU 3</td><td>10X v2</td><td>8000</td><td>4067</td><td>3125</td><td>3103</td><td>441</td><td>2598</td><td colspan="2">51</td><td>18</td></tr>
<tr><td>LF</td><td>10X v2</td><td>8000</td><td>2383</td><td>4938</td><td>2541</td><td>864</td><td>118</td><td>39</td><td rowspan="4">60</td><td>40</td></tr>
<tr><td>RF</td><td>10X v2</td><td>8000</td><td>2697</td><td>5102</td><td>2622</td><td>848</td><td>123</td><td>37</td><td>48</td></tr>
<tr><td>LH</td><td>10X v2</td><td>8000</td><td>2122</td><td>3755</td><td>2711</td><td>598</td><td>183</td><td>28</td><td>40</td></tr>
<tr><td>RH</td><td>10X v2</td><td>8000</td><td>1095</td><td>5261</td><td>2354</td><td>789</td><td>138</td><td>35</td><td>40</td></tr>
</table>

**Supplementary Table 4: Statistical significance (p-value) of differences in average HSC-rooted clone size (Figure 6)**

| … has Larger Clones than… | | Sample 2? | | | |
|---|---|---|---|---|---|
| | | **5FU Mouse 1** | **Control 1** | **Control 2** | **Control 3** |
| **Sample 1** | **5FU Mouse 1** | 1.0e+00 | 1.7e-04 | 1.1e-07 | 7.2e-11 |
| | **Control 1** | 1.0e+00 | 1.0e+00 | 9.5e-01 | 3.7e-01 |
| | **Control 2** | 1.0e+00 | 1.0e+00 | 1.0e+00 | 5.1e-01 |
| | **Control 3** | 1.0e+00 | 1.0e+00 | 1.0e+00 | 1.0e+00 |

**Supplementary Table 5: Differential gene expression between parent HSCs and childless HSCs (Figure 6)**

| Gene | p_val | avg_logFC | pct.1 | pct.2 | p_val_adj |
|---|---|---|---|---|---|
| *Plac8* | 1.11E-05 | 0.443686 | 1 | 1 | 0.022181 |
| *Cdk6* | 0.000146 | 0.30155 | 0.971 | 0.988 | 0.292948 |
| *Nkg7* | 0.000342 | 0.455855 | 0.829 | 0.554 | 0.683012 |
| *Phgdh* | 0.0005 | 0.28048 | 1 | 0.926 | 1 |
| *Emb* | 0.000648 | 0.335612 | 0.943 | 0.678 | 1 |
| *Smarca2* | 0.002105 | -0.29964 | 0.629 | 0.769 | 1 |
| *Ablim1* | 0.003223 | 0.26776 | 0.714 | 0.459 | 1 |
| *Adssl1* | 0.003399 | 0.2906 | 0.857 | 0.649 | 1 |
| *Pnisr* | 0.004244 | -0.30633 | 0.971 | 0.988 | 1 |
| *Serpinb1a* | 0.004474 | 0.282037 | 0.914 | 0.793 | 1 |
| *Cd81* | 0.005061 | -0.28216 | 0.943 | 0.971 | 1 |
| *Muc13* | 0.005277 | 0.255475 | 0.886 | 0.583 | 1 |
| *Stat1* | 0.005635 | 0.30276 | 0.771 | 0.541 | 1 |
| *Cd24a* | 0.011899 | -0.31708 | 0.8 | 0.781 | 1 |
| *Mpo* | 0.017957 | 0.322474 | 1 | 0.992 | 1 |
| *Taok3* | 0.027004 | 0.285475 | 0.8 | 0.674 | 1 |
| *Glul* | 0.04018 | -0.33248 | 0.857 | 0.793 | 1 |
| *Cd48* | 0.063211 | 0.435107 | 0.6 | 0.488 | 1 |
| *Cpne2* | 0.09082 | 0.265093 | 0.829 | 0.665 | 1 |
| *Tbxas1* | 0.098405 | -0.2673 | 0.771 | 0.711 | 1 |
| *Plxnc1* | 0.099949 | -0.29844 | 0.6 | 0.574 | 1 |
| *Slc22a3* | 0.118351 | 0.257604 | 0.571 | 0.421 | 1 |
| *Bex4* | 0.119088 | -0.29402 | 0.6 | 0.579 | 1 |
| *Shisa5* | 0.130016 | -0.27373 | 1 | 0.893 | 1 |
| *Pf4* | 0.314527 | -0.26528 | 0.629 | 0.612 | 1 |
| *Meg3* | 0.329864 | -0.31519 | 0.886 | 0.74 | 1 |
| *Adgrg3* | 0.395805 | 0.264516 | 0.686 | 0.583 | 1 |
| *Mllt3* | 0.399855 | -0.27227 | 0.8 | 0.657 | 1 |
| *Ltb* | 0.41547 | -0.27331 | 0.857 | 0.781 | 1 |
| Mycn | 0.964807 | -0.26505 | 0.686 | 0.537 | 1 |

**Supplementary Table 6: Differential gene expression between parent and childless HSC cluster (Figure 6)**

| Gene | p_val | avg_logFC | pct.1 | pct.2 | p_val_adj |
|---|---|---|---|---|---|
| *Plac8* | 2.71E-175 | 0.546865 | 0.998 | 0.988 | 5.42E-172 |
| *Cd34* | 1.37E-124 | 0.412811 | 0.957 | 0.896 | 2.74E-121 |
| *Cdk6* | 7.55E-103 | 0.409562 | 0.979 | 0.965 | 1.51E-99 |
| *H2afy* | 3.52E-137 | 0.374018 | 1 | 0.998 | 7.04E-134 |
| *Serpinb1a* | 1.80E-84 | 0.369092 | 0.859 | 0.786 | 3.60E-81 |
| *Mpo* | 4.50E-56 | 0.36604 | 0.971 | 0.939 | 9.01E-53 |
| *Nkg7* | 6.63E-55 | 0.34021 | 0.64 | 0.51 | 1.33E-51 |
| *Fam117a* | 1.91E-39 | 0.278728 | 0.673 | 0.582 | 3.82E-36 |
| *Sox4* | 4.64E-47 | 0.252392 | 0.996 | 0.992 | 9.28E-44 |
| *Mecom* | 3.53E-34 | -0.25409 | 0.596 | 0.751 | 7.07E-31 |
| *Jund* | 9.00E-35 | -0.25649 | 0.878 | 0.948 | 1.80E-31 |
| *Fos* | 9.94E-18 | -0.259 | 0.609 | 0.73 | 1.99E-14 |
| *Mmrn1* | 2.55E-32 | -0.26561 | 0.557 | 0.669 | 5.09E-29 |
| *Jun* | 6.71E-29 | -0.2684 | 0.894 | 0.965 | 1.34E-25 |
| *Rgs1* | 3.31E-15 | -0.27309 | 0.648 | 0.728 | 6.62E-12 |
| *Junb* | 2.32E-35 | -0.2844 | 0.59 | 0.754 | 4.64E-32 |
| *Ltb* | 3.87E-30 | -0.28618 | 0.758 | 0.845 | 7.74E-27 |
| *Ier2* | 1.75E-45 | -0.30564 | 0.786 | 0.904 | 3.50E-42 |
| *Egr1* | 1.20E-29 | -0.31212 | 0.495 | 0.676 | 2.39E-26 |
| *Pdzk1ip1* | 3.38E-58 | -0.3234 | 0.641 | 0.808 | 6.75E-55 |
| *Tbxas1* | 2.06E-77 | -0.33632 | 0.596 | 0.809 | 4.13E-74 |
| *Hacd4* | 8.54E-65 | -0.3538 | 0.628 | 0.799 | 1.71E-61 |
| *Neat1* | 1.04E-56 | -0.36253 | 0.818 | 0.926 | 2.07E-53 |
| *Apoe* | 5.11E-63 | -0.42971 | 0.582 | 0.782 | 1.02E-59 |
| *Mllt3* | 9.18E-67 | -0.44728 | 0.514 | 0.717 | 1.84E-63 |
| *Meg3* | 1.08E-130 | -0.55864 | 0.486 | 0.728 | 2.16E-127 |
| *Jchain* | 1.08E-10 | -0.64067 | 0.503 | 0.624 | 2.17E-07 |
| *Ighm* | 0.000106 | -0.82665 | 0.517 | 0.575 | 0.211726 |
